# Memory Trace Imbalance in Reinforcement and Punishment Systems Can Reinforce Implicit Choices Leading to Obsessive-Compulsive Behavior

**DOI:** 10.1101/2020.08.07.241588

**Authors:** Yuki Sakai, Yutaka Sakai, Yoshinari Abe, Jin Narumoto, Saori C. Tanaka

## Abstract

We may view most of our daily activities as rational action selections; however, we sometimes reinforce maladaptive behaviors despite having explicit environmental knowledge. In this study, we model obsessive-compulsive disorder (OCD) symptoms as implicitly learned maladaptive behaviors. Simulations in the reinforcement learning framework show that agents implicitly learn to respond to intrusive thoughts when the memory trace signal for past actions decays differently for positive and negative prediction errors. Moreover, this model extends our understanding of therapeutic effects of behavioral therapy in OCD. Using empirical data, we confirm that patients with OCD show extremely imbalanced traces, which are normalized by serotonin enhancers. We find that healthy participants also vary in their obsessive-compulsive tendencies, consistent with the degree of imbalanced traces. These behavioral characteristics can be generalized to variations in the healthy population beyond the spectrum of clinical phenotypes.

## Introduction

We live in a complex and dynamic environment and choose better behaviors based upon trial and error. Although we may view most of our daily activities as rational action selections based on our environmental knowledge, in reality, this is not always the case. For example, we may repeatedly study hard even if we are already well prepared for an exam. We may also find ourselves abiding by superstitions, even though we know they make no sense. In these situations, there is a minor discrepancy between explicit environmental understanding, i.e., action-outcome knowledge, and implicit learning. Learning is implicit when people have become sensitive to regularities of the environment in the absence of declarative knowledge (Cleeremans, 2009). In cases in which these discrepancies between explicit environmental understanding and implicit learning are significant, they are accompanied by “ego-dystonic” anxiety, which is a characteristic of obsessive-compulsive disorder (OCD) (Gillan and Robbins, 2014; Vaghi et al., 2017). In patients with OCD, “ego-dystonic” anxiety becomes lodged in their minds (Salkovskis, 1999). To neutralize and relieve the anxiety, they tend to perform compulsive actions, despite their cost, e.g., excessively washing their hands or repeatedly checking their keys, even though they know that these behaviors are excessive and unreasonable (Cavedini et al., 2006).

We must consider theoretical and biological implementations when investigating implicit learning leading to ego-dystonic anxiety. Implicit trial and error learning requires repetitive experiences, and reinforcement learning theory provides a framework for learning appropriate choices through repetitive experiences (Sutton and Barto, 1998). Reinforcement learning enhances or punishes recent choices based on differences between actual outcomes and predictions, called prediction errors. Positive prediction errors reinforce recent choices, whereas negative errors punish them.

When examining biological implementations of implicit learning, it is essential to consider previous work demonstrating that the activity of dopamine-projecting neurons in the midbrain resembles prediction errors (Schultz et al., 1997). Dopaminergic neurons project widely to cortical and subcortical areas. One of the main targets of dopamine is the striatum, which is located in the basal ganglia (Graybiel, 2000). In the striatum, distinct types of dopamine receptors—D1 and D2 receptors—are expressed in exclusive neuron groups, which are thought to be involved in distinct circuit patterns, the direct and indirect pathways (Graybiel, 2000). Knowledge of these pathways has accumulated primarily about initiation and termination of motor actions and about selecting task-relevant cues. However, some studies suggested that these pathways optimize action selection based on its outcome in a pathway-specific manner (Kravitz et al., 2012; Nonomura et al., 2018). Artificial activation of D1- and D2-expressing neurons after a certain behavioral action reinforces and punishes the action, respectively (Kravitz *et al*., 2012). Direct and indirect pathway neurons respond to positive and negative outcomes, respectively (Nonomura *et al*., 2018). These studies imply that reinforcement and punishment of recent actions may be reflected in the direct and indirect pathways, respectively.

In implementation of implicit learning, we should also consider delays as well as reinforcement/punishment. In general, the outcome of a certain choice is available after a certain delay. Therefore, reinforcement and punishment should be assigned to recent choices within a certain time scale. This is called credit assignment, which is implemented as eligibility traces in reinforcement learning (Sutton and Barto, 1998). An eligibility trace can be implemented as a memory trace in each synapse (Gerstner et al., 2018; Izhikevich, 2007; Yagishita et al., 2014). In such case, trace time scales for reinforcement and punishment should be separately controlled in direct and indirect pathways. Reinforcement learning theory requires that both trace time scales be equal. However, this requirement cannot be completely realized in distinct neural systems.

Since abundant evidence suggests that an imbalance between direct and indirect pathways is central to the pathophysiology of OCD (Abe et al., 2015; Pauls et al., 2014; Sakai et al., 2011), we hypothesized that obsessive-compulsive behaviors are generated by imbalanced implicit learning between reinforcement and punishment. Furthermore, our previous study reported modulatory effects of the balance of trace time scales by serotonin (Tanaka et al., 2009), while serotonin reuptake inhibitors (SRIs) show the therapeutic effects in OCD (Koran et al., 2007). Given these two factors, it is plausible that an altered balance of trace time scales between reinforcement and punishment might be the core mechanism of OCD. However, it is unclear how the balance of trace time scales leads to development of OCD and whether other learning parameters can explain it. In addition, previous OCD models can explain extreme reinforcement of compulsion (repetitive behaviors to temporarily relieve the anxiety caused by obsession), but they cannot explain the increased frequency of obsession (recurrent and persistent thoughts that cause anxiety) (Banca et al., 2014; Gillan and Robbins, 2014; Hauser et al., 2017). Therefore, we combined computational theory and experimental validation to identify its mechanism.

First, we propose a computational model that enables implementation of imbalanced learning between reinforcement and punishment. We conducted a simulation showing that imbalanced learning reinforces implicit learning, leading to a spiral of repetitive obsession and compulsion, even if agents have intact environmental understanding. Second, we tested our hypothesis in a behavioral task for healthy participants and for participants with OCD. In addition, we evaluated mechanisms of first-line treatments, i.e., behavioral therapy and psychotropic medication for OCD, in our computational model and behavioral data. Finally, we investigated whether these behavioral characteristics could be generalized to variations in the healthy population, beyond the spectrum of clinical phenotypes.

## Results

### Theoretical model of OCD

We described mental states involved in obsessive-compulsive symptoms in stochastic transitions between two states: a relief state and an obsessive state (Figure 1a). We assumed an action to relieve anxiety as an option in the obsessive state, which could stochastically permit a transition from the obsessive state to the relief state, at a certain cost. Such an action might become compulsive in OCD, and we labeled it as a “compulsion” (Figure 1a). We combined any other options in the obsessive state, which could relieve anxiety with lower probability without any cost into the option “other” (Figure 1a). Even in the relief state, even healthy people sometimes experience idle thoughts called intrusive thoughts (Salkovskis, 1999). Therefore, an essential difference between patients with OCD and healthy people is whether they reinforce an abnormal reaction to intrusive thoughts (Salkovskis, 1999). Accordingly, we labeled the transition from the relief state to the obsessive state as an “abnormal reaction to intrusive thoughts” mainly seen in OCD (Salkovskis, 1999) (Figure 1a). We also combined other options that maintain the relief state into “other” (Figure 1a). We assumed that every stay in the obsessive state would produce a negative outcome representing anxiety. No positive outcome was introduced in the state transition diagram imitating obsessive compulsive symptoms (Figure 1a). Hence, maintaining “other” in the relief state is clearly optimal. Normal reinforcement learning should reduce abnormal reactions to intrusive thoughts. However, if the learning system has some defects, it may fail to reduce the reaction to intrusive thoughts and may fall into a spiral of obsession and compulsion. This is the discrepancy between explicit environmental understanding and implicit learning, since agents fall into a spiral of obsession and compulsion even though they know that they will experience only loss.

**Figure 1.**
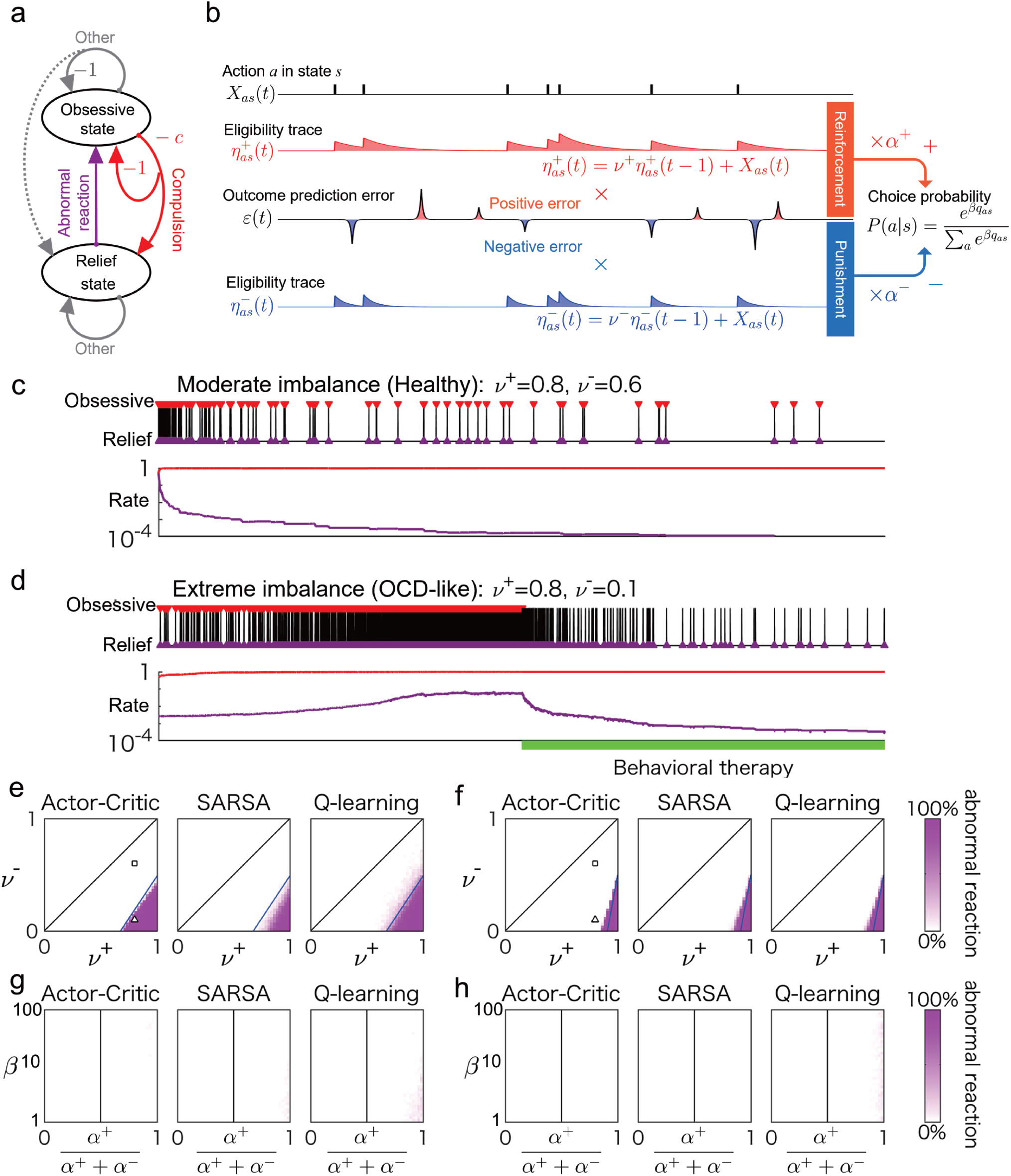
OCD-like behavior caused by imbalanced implicit learning. (a) State transition diagram imitating obsessive compulsive symptoms. The dashed arrow means that a low transition probability was supposed. Outcomes were zero in transitions without descriptions of outcomes. (b) Schema of the separate eligibility trace model. (c) Simulation of the separate eligibility trace model in the state transition diagram with a moderate imbalance in trace factors, *ν*^±^. Black polylines represent the temporal pattern of state transitions. Red and magenta triangles represent occurrences of compulsions and abnormal reactions to intrusive thoughts, respectively. The lower plots show temporal patterns of compulsion (red curve) and abnormal-reaction (magenta curve) rates on a logarithmic scale. The upper plots show the simulated results of occurrences of actions or state transitions, while the lower plots represent the variables of the computational model. (d) As in (c), but in the case of extreme imbalance. In the latter half, we demonstrated behavioral therapy by preventing the compulsion in the obsessive state (green bar). (e) Conditions of OCD-like behavior in the trace factor space, *ν*^±^ with different types of learning algorithms: actor-critic, SARSA, and Q-learning. The magenta color scale represents the percentage increase of the abnormal-reaction rate among 10 simulations around optimal behavior (zero reaction rate to intrusive thoughts). Solid blue lines represent the theoretically derived boundary (see the theoretical derivation section of STAR Methods). The square and triangle represent the trace factors *ν*^±^ used in (c) and (d). (f) As in (e), but in this condition, behavioral therapy did not work. (g)(h) As in (e) and (f), but the learning-rate imbalance *α*^+^/(*α*^+^ + *α*^−^) and the greediness parameter *β* were swept under the balanced trace factors *ν*^+^ = *ν*^−^. The grid for *β* is in logarithmic scale.

We implemented imbalanced learning with a reinforcement learning model separated into reinforcement and punishment systems, which we call *a separate eligibility trace model* (see STAR Methods). Each trace represents the recent frequency of action in a specific state within a time scale determined by the trace decay factor, *ν*. If eligibility traces for positive and negative prediction errors are implemented in distinct neural systems, the respective eligibility traces 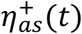 and 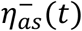 at time step *t* for action *a* (*a* = 0, 1, …) in state *s* (*s* = 0, 1, …) obey the following equation (Sutton and Barto, 1998),

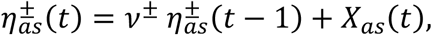

where *X*_*as*_ (*t*) = 1 when action *a* is chosen in state *s* at time step *t* and *X*_*as*_ (*t*) = 0 otherwise. When a prediction error occurs, the choice probability of each action in each state is updated by the product of the eligibility trace and the prediction error (Figure 1b). Thus, a positive prediction error reinforces recent actions within a time scale, whereas a negative error punishes them. Theoretically, trace factors in the two systems, *ν*^±^, should be balanced: *ν*^+^ = *ν*^−^. However, separate neural implementations make it difficult to maintain the balance perfectly. In our separate eligibility trace model, we assumed that an imbalance in the trace factors can occur: *ν*^+^ ≠ *ν*^−^.

To show whether imbalanced learning generates OCD-like behavior, we simulated actor-critic learning (STAR Methods), a typical method of reinforcement learning models (Sutton and Barto, 1998) in the state transition diagram (Figure 1a), incorporating a separate eligibility trace (Figure 1b). When the imbalance in the trace factors was moderate, actor-critic learning reduced the reaction rate to intrusive thoughts, even when the simulation started with a high reaction rate to intrusive thoughts (Figure 1c). In contrast, extreme imbalance increased the reaction rate to intrusive thoughts such that it culminated in a spiral of repetitive obsession and compulsion (Figure 1d). Such OCD-like behavior was reduced by preventing the compulsion during the obsessive state (Figure 1d). This is the behavioral therapy of exposure and response prevention (ERP) and is one of the first-line treatments for OCD (Koran *et al*., 2007). ERP is the therapy to break the spiral of obsession and compulsion by helping patients how to confront anxiety and select actions other than compulsions.

Because the situation has no positive outcomes, positive prediction errors can occur only in the state transition from the obsessive state (negative value state) back to the relief state (near zero-value state). At that time, the positive eligibility trace reinforces the recent abnormal reaction in a relief state, if the positive trace factor *ν*^+^ is large enough. On the other hand, negative outcomes during a stay in the obsessive state cannot sufficiently punish the abnormal reaction if the negative trace factor *ν*^−^ is small. When *ν*^±^ is balanced, the effect of punishment on the abnormal response will always be more significant than reinforcement. However, if the agent shows excessive imbalance *ν*^+^ > *ν*^−^, the abnormal reaction would be reinforced since the dominance of reinforcement and punishment would be inverted. That is the mechanism of OCD-like behavior by trace-factor imbalance.

We identified the condition for the trace factor *ν*^±^ in which OCD-like behavior might emerge by numerical simulations in a representative set of other model parameters (Figure 1e; STAR Methods). In addition to actor-critic learning (Figure 1e), we simulated SARSA (State-Action-Reward-State-Action) and Q-learning (Figure 1e), other typical reinforcement learning models (Sutton and Barto, 1998). The color map represents the fraction of 100 simulation runs in which abnormal reactions to intrusive thoughts were reinforced at each pair of (*ν*^+^, *ν*^−^). We identified conditions of OCD-like behavior in the region *ν*^+^ > *ν*^−^ (Figure 1e). In such cases, the trace scale for reinforcement was longer than that for punishment. We also identified conditions in which behavioral therapy of ERP would not work and localized it to a region with a more robust imbalance (Figure 1f).

In the downstream of the separate eligibility traces, the learning rate *α* is multiplied (Figure 1b). Therefore, we can suppose an imbalance of the learning rate. After the separate streams are merged through the update of each policy parameter *q*_*as*_, the greediness parameter *β* controls the soft-max choice probabilities to determine the exploration and exploitation balance (see STAR Methods). The imbalance in the learning rate, *α*, has often been discussed in different contexts from OCD (Lefebvre et al., 2017; Palminteri and Lebreton, 2022), and an excessively large greediness parameter, *β*, has been discussed in relation to persistence of compulsion. Hence, we confirmed whether OCD-like behavior emerges because of the imbalance in the learning rate, *α*, or an excessively large *β* in the same situation shown in Figures 1e and 1f (see STAR Methods). Except for accidental reinforcements of abnormal reactions to intrusive thoughts, steady abnormal reactions did not emerge in any bias of the learning-rate imbalance *α*^+^/(*α*^+^ + *α*^−^) and larger *β* up to 10 (Figures 1g and 1h), when the trace factors were balanced *ν*^+^ = *ν*^−^.

We derived the theoretical conditions as functions of other parameters in the three learning models: actor-critic, SARSA, and Q-learning (see the theoretical derivation section of STAR Methods). In each learning model, we found that the reinforcement of the abnormal reaction requires a necessary condition *ν*^+^ > *ν*^−^. This result suggests that a person with *ν*^+^ > *ν*^−^ has a risk of OCD. The imbalance in learning rate α modulates the condition, but it can never cause reinforcement of an abnormal reaction with balanced trace factors *ν*^+^ = *ν*^−^. The greediness parameter *β* is irrelevant to the condition. Thus, the trace-factor imbalance *ν*^+^ > *ν*^−^ can potentially cause OCD-like behavior, whereas an imbalance in learning rate *α* or a large greediness parameter *β* alone can never cause it.

Each OCD patient presents a certain compulsive action for specific anxieties among many types of anxiety. The cost of compulsive action and the probability of relieving the anxiety depend on the type of anxiety. Therefore, corresponding parameters of the state transition diagram (Figure 1a) should differ by the kind of anxiety. Next, we considered whether a certain anxiety situation to present OCD-like behavior exists among any type of anxiety situation. Namely, we derived the condition in which the state transition parameters (Figure 1a) exist such that abnormal reactions would be reinforced. We found the common condition that *α*^+^(*ν*^+^−*ν*^−^) > *α*^−^*γ* in actor-critic, SARSA, and Q-learning (see the theoretical derivation section of STAR Methods), where the discount factor *γ* ≥ 0 is a parameter in the reward prediction error. We can see in the inequality that the learning-rate imbalance just modulates the condition with the trace-factor imbalance.

### Empirical evaluation of the theoretical model of OCD

To empirically test the suggestion that *ν*^+^ > *ν*^−^, derived from our computational model of OCD, we subjected patients with OCD (n = 45) and healthy controls (HCs) (n = 168) to a behavioral task. We excluded 3 OCDs due to Intelligence Quotient (IQ) < 80 using the Wechsler Adult Intelligence Scale - Third Edition (WAIS-III). Seventeen HCs were excluded for medical or experimental reasons. The subsequent analysis was conducted with 42 patients with OCD and 151 HCs. See STAR Methods and Supplementary Table 1 for detailed information about participants.

We used a delayed feedback task to evaluate trace factors *ν*^±^, as in our previous research (Tanaka *et al*., 2009), except for presented stimuli (abstract cues). Briefly, participants chose one of two options displayed on a monitor by pressing a left or right button in each trial (Figure 2a). Each stimulus represented different outcomes (+10, +40, −10, or −40 yen) and a delay to the feedback (immediately after a button press or three trials later) (Figure 2b and c). For example, +40(0) represented a gain of 40 yen during the current trial (immediate reward), and −10(3) represented a loss of 10 yen after three trials (delayed punishment). Participants were not told about stimulus-outcome associations (Figure 2b). They received money after the experiment in proportion to the total outcome obtained in the delayed feedback task. To maximize the total outcome, participants needed to learn appropriate stimulus choices by correctly assigning the credit of the current feedback to the recent choices that caused the current feedback. The difference in learning effects between immediate and delayed feedback reflected the trace factors, *ν*^±^.

**Figure 2.**
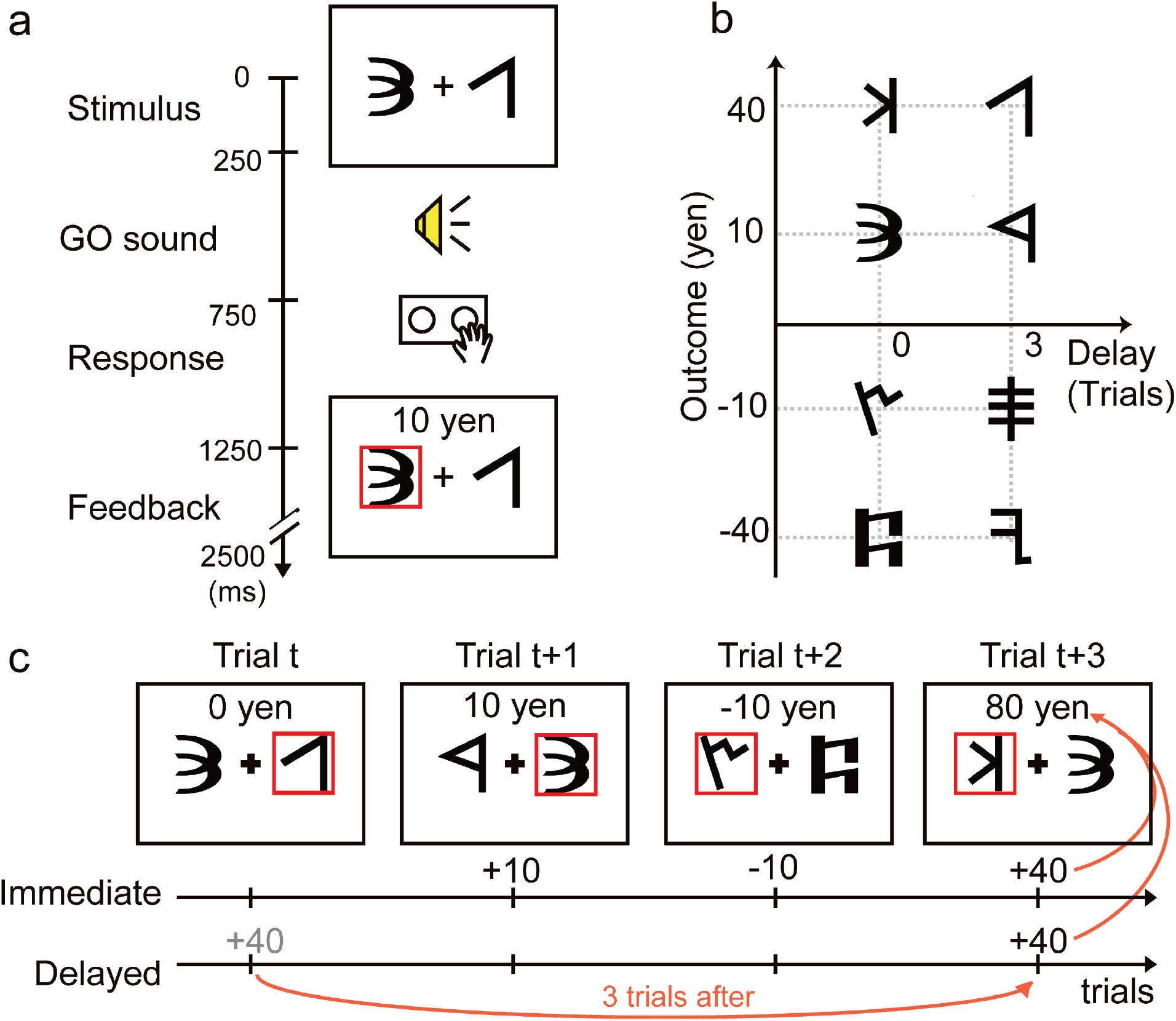
Delayed feedback task. (a) Two abstract cues were displayed in each trial. When participants heard an auditory cue (beep), they chose one of the stimuli within 1 s. A single trial lasted 2.5 s. Sequences of stimulus pairs were pseudorandom. (b) Outcome-delay mapping for each stimulus. Mapping differed among participants. (c) An example of a delayed feedback task. If participants chose the stimulus with a delay [+40(3)] at trial *t*, that outcome was not displayed immediately. If they chose the stimulus with no delay [+40(0)] at trial *t*+3, the sum of the delayed and immediate outcomes was displayed at trial *t*+3 (in this case +80 yen).

Based on our simulation of OCD-like behavior with trace factors *ν*^+^ > *ν*^−^ (Figure 1e), patients with OCD showed impaired learning involving stimuli with delayed feedback. To confirm relationships between the parameters (*ν*^+^ and *ν*^−^) and the total outcome from stimuli with immediate or delayed feedback, we conducted a simulation (STAR Methods). The total outcome from stimuli with immediate feedback did not depend on the parameters, *ν*^+^ and *ν*^−^, except for extremely high values (Figure 3a). In contrast, we confirmed that more balanced and higher values of *ν*^+^ and *ν*^−^ were needed to achieve a greater total outcome from stimuli with delayed feedback (Figure 3a). Therefore, the total outcome from stimuli with immediate or delayed feedback was compared between groups.

**Figure 3.**
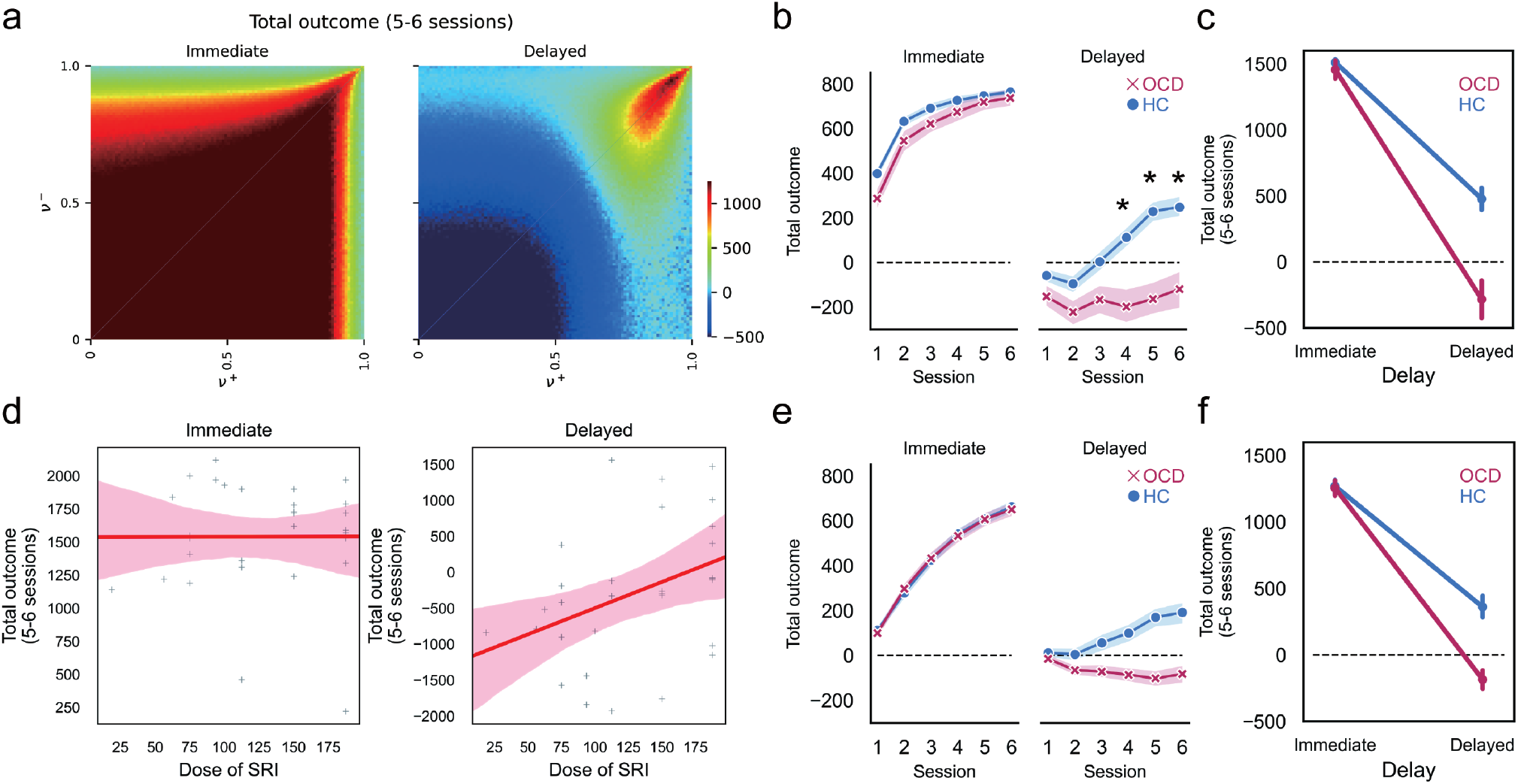
Learning of stimuli with immediate or delayed feedback and its relationship with the dose equivalence of SRIs. (a) The total simulated outcome from stimuli with immediate (left) or delayed feedback (right) projected in *ν*^+^/*ν*^−^ space. The color represents the amount of total outcome in sessions 5 and 6. (b) The total outcome from stimuli with immediate (left) or delayed feedback (right) (red cross, patients with OCD; blue circle, HCs). The line and colored area represent the mean and standard error. The horizontal dashed line at 0 yen. *p_adjusted_ < 0.05 with Bonferroni-Holm–corrected post-hoc comparisons of the simple main effects of group (patients with OCD < HCs). (c) Significant interactions between delays (immediate and delayed) and groups (red, patients with OCD; blue, HCs) in the total outcome in sessions 5 and 6. The line and error bar represent the mean and standard error. The horizontal dashed line at 0 yen. (d) Correlation between the SRI dose and the total outcome in sessions 5 and 6 (left, immediate feedback; right, delayed feedback). The line and colored areas are the regression line and the 95% bootstrapped confidence interval (+: each OCD patient taking SRIs). (e) As in (b), but in the case of simulated data using estimated parameters (red cross, simulated patients with OCD; blue circle, simulated HCs). (f) As in (c), but in the case of simulated data using estimated parameters (red, simulated patients with OCD; blue, simulated HCs).

Consistent with our trace factor hypothesis, the OCD group showed impaired learning of stimuli with delays (Figure 3b). We used the ANOVA-type statistic (ATS) for nonparametric repeated measures ANOVA (Noguchi et al., 2012) with a within-participant factor of sessions (1–6 sessions) and a between-participant factor of groups (OCD and HC). There were significant interactions in learning stimuli with delays (F(2.71, ∞)=5.35, p=0.0017). Bonferroni-Holm– corrected post-hoc comparisons using the Brunner-Munzel test confirmed OCD < HC in session 4 (statistic = −3.72, p_adjusted_ = 0.0017), session 5 (statistic = −4.93, p_adjusted_ < 0.001), and session 6 (statistic = −3.94, p_adjusted_ < 0.001) (Figure 3b). In contrast, patients with OCD showed intact learning of stimuli with immediate feedback (Figure 3b). There were no significant interactions (F(3.01, ∞)=1.04, p > 0.05) and there was no main effect of groups (F(1, 56.5)=2.57, p > 0.05). To better clarify the effect of delay on learning, we conducted an ATS analysis of a within-participant factor of delays (immediate and delayed) and a between-participant factor of groups using the total outcome in sessions 5 and 6. We confirmed that there were significant interactions (F(1, ∞)=5.17, p=0.023). That is, a delay in receiving feedback impaired learning more in patients with OCD than in HC (Figure 3c).

Considering the therapeutic effect of SRIs in OCD (Koran *et al*., 2007), SRIs may normalize impaired learning of stimuli with delayed feedback. To investigate effects of SRIs on learning, we conducted a correlation analysis between the dose equivalence of SRIs (Inada and Inagaki, 2015) (STAR Methods and Supplementary Table 2) and the total outcome from stimuli with immediate or delayed feedback in sessions 5 and 6. Consistent with our prediction, SRI dose was significantly correlated with the total outcome from stimuli with delayed feedback [Spearman’s rank correlation (27); r = 0.41, p = 0.027; Figure 3d], but not with immediate feedback [Spearman’s rank correlation (27); r = 0.061, p > 0.05; Figure 3d]. Since there was no significant correlation between total outcome and depressive symptoms evaluated using the 17-item Hamilton Depression Rating Scale (HDRS) (Hamilton, 1967) [Immediate: Spearman’s rank correlation (26); r = 0.14, p > 0.05; Delayed: Spearman’s rank correlation (26); r = −0.20, p > 0.05], a simple antidepressant effect of SRIs could not explain this improved learning caused by SRIs.

In summary, patients with OCD demonstrated the hypothesized impaired learning involving stimuli with delayed feedback and SRIs improved it in a dose-dependent manner.

### Estimation of model parameters from empirical data

Even though the results of total outcome (Figure 3b) suggested that HC may have more balanced and higher values of *ν*^+^ and *ν*^−^ than OCD (Figure 3a), it is difficult to use this as direct evidence of the trace-factor imbalance *ν*^+^ > *ν*^−^ hypothesized in OCD. To investigate direct evidence of the trace-factor imbalance *ν*^+^ > *ν*^−^ in OCD, we fitted behavioral data to the actor-critic learning model using the learning rate (*α*), exploration-exploitation degree (*β*), and separation of the eligibility traces for reinforcement and punishment (*ν*^±^) (STAR Methods). Because we were specifically interested in the individual variance represented by reinforcement learning parameters, we fitted parameters in data from each participant independently, using maximum a posteriori estimation rather than pooled data of a group (Daw, 2011).

We simulated behavioral data of each group with the estimated parameters to validate the actor-critic learning model with separate eligibility traces (see STAR Methods). We evaluated whether all-important behavioral characteristics are qualitatively and quantitatively captured (Wilson and Collins, 2019). We confirmed that the specific impairment in learning from stimuli with delayed feedback in the simulated OCD group (Figure 3e and 3f) resembles the experimental data (Figure 3b and 3c). Therefore, our model actually captured essential features of human behaviors in this task.

To clarify the *ν*^+^/*ν*^−^ distribution of each group, we projected estimated parameters from each participant to *ν*^+^/*ν*^−^ space and visualized them using kernel density estimation (Figure 4a). Consistent with our computational model simulation of OCD, patients with OCD were distributed in the *ν*^+^ > *ν*^−^ imbalanced area, whereas the distribution was balanced in HCs (Figure 4a). To compare the *ν*^+^/*ν*^−^ distribution among groups, we confirmed the multivariate homogeneity of group dispersions [F (1, 191) = 0.25, p = 0.62] and applied permutational multivariate analysis of variance (PERMANOVA). The *ν*^+^/*ν*^−^ distribution differed significantly between groups [PERMANOVA: F (1, 191) = 5.99, p = 0.0060] (Figure 4a). Parameters *α* and *β* did not differ significantly between groups (Brunner-Munzel test, p > 0.05). Among OCD participants taking SRIs (n = 29), the equivalent dose of SRIs was significantly correlated with the imbalanced settings of *ν* (= *ν*^+^ − *ν*^−^) [Spearman’s rank correlation (27); r = −0.48, p = 0.086; Figure 4b]. That is, higher doses of SRIs normalized the imbalanced settings of *ν* in participants with OCD.

**Figure 4.**
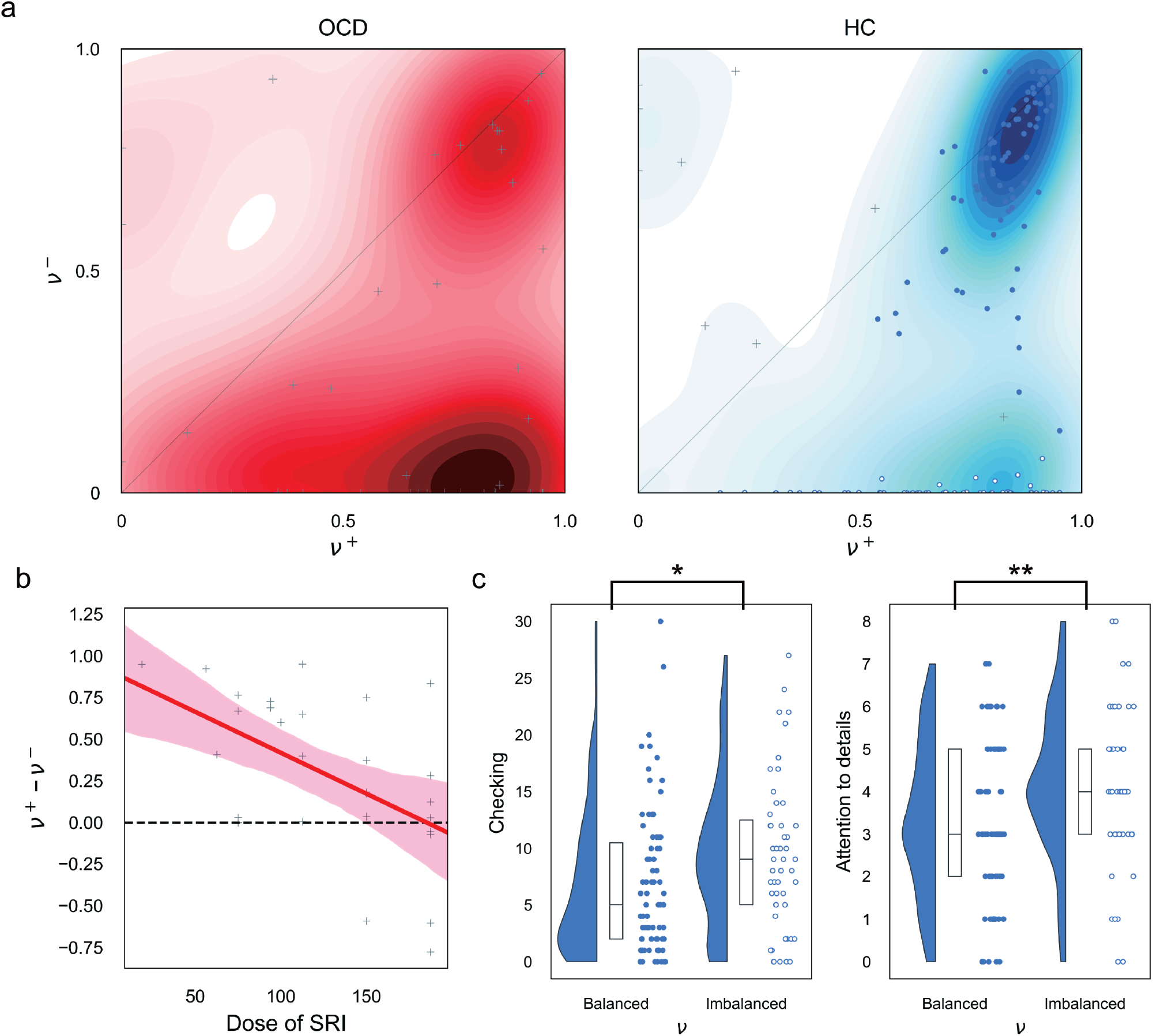
Estimated parameters and their relationships with clinical characteristics. (a) Distribution of participants in each group projected in *ν*^+^/*ν*^−^ space using the kernel density estimation (red, patients with OCD; blue, HCs). The horizontal axis represents *ν*^+^ and the vertical axis represents *ν*^−^. The diagonal line represents the balanced line in *ν*^+^/*ν*^−^ space. Blue and white circles in the right figure represent balanced and imbalanced clusters in HCs, respectively. Other participants were depicted using +. (b) Significant correlation between the SRI dose and *ν*^+^ − *ν*^−^ [Spearman’s rank correlation (27); r = −0.48, p = 0.0086]. The line and colored areas are the regression line and the 95% bootstrapped confidence interval (+: each OCD patient taking SRIs). The dashed line represents the balanced settings of *ν*^±^. (c) PI Checking and AQ Attention to Detail scores in the balanced and imbalanced clusters in HCs. The half-violin, box, and strip plots represent the probability density, interquartile range and median, and raw data, respectively. Blue and white circles represent balanced and imbalanced clusters, respectively, whereas * and ** represent significant differences (Brunner-Munzel test, * statistic = −2.28, p = 0.012; ** statistic = −2.82, p = 0.0028).

Although the *ν*^+^/*ν*^−^ distribution was more balanced in HCs compared with patients with OCD, we found some diversity among HCs. We conducted clustering analysis using hierarchical density-based spatial clustering of applications with noise (HDBSCAN) (Campello et al., 2015) to evaluate diversity among HCs. HDBSCAN revealed two clusters of HCs: a balanced *ν* cluster (Figure 4a, blue circle; n = 83) and an imbalanced *ν* cluster (Figure 4a, white circle; n = 57), with 11 HCs that were not clustered. Obsessive-compulsive trends and a propensity to adhere to fine-grained details were compared between clusters using five subscales of the Padua Inventory (PI; “Checking”, “Dirt”, “Doubt”, “Impulse”, “Precision”) (Sugiura and Tanno, 2000) and the Attention to Detail subscale of the Autism Quotient (AQ) (Wakabayashi et al., 2004), respectively. The imbalanced HC cluster showed significantly higher PI Checking scores (Brunner-Munzel test, statistic = −2.28, p = 0.012), PI Dirt scores (Brunner-Munzel test, statistic = −1.71, p = 0.045), and AQ Attention to Detail scores (Brunner-Munzel test, statistic = −2.82, p = 0.0028) than the balanced HC cluster. Significant differences were still found in the PI Checking score and the AQ Attention to Detail score after false discovery rate correction for multiple comparisons among scales (Figure 4c) (PI Checking, p_adjusted_ = 0.025; AQ Attention to Detail, p_adjusted_ = 0.01).

## Discussion

This study showed that imbalanced implicit learning between reinforcement and punishment can induce a spiral of repetitive obsession and compulsion, even despite explicit environmental understanding. We constructed a computational model of OCD using a separate eligibility trace model (Figure 1a-e) and found extremely imbalanced trace factors *ν*^+^ > *ν*^−^ in OCD (Figure 4a). In addition, effects of behavioral therapy (ERP) and psychotropic medication (SRIs), which are the first-line treatments for OCD, were reflected in our computational model (Figure 1d) and behavioral results (Figure 3d and 4b), respectively.

While the theoretical framework of the eligibility trace has long been conceptualized in the field of reinforcement learning (Sutton and Barto, 1998), empirical evidence for it has been reported only relatively recently (Brzosko et al., 2015; Brzosko et al., 2017; He et al., 2015; Yagishita *et al*., 2014). The latest theories regarding synaptic plasticity have proposed that co-activation of pre- and postsynaptic neurons sets a flag at the synapse (eligibility trace) that leads to a weight change only if additional factors, i.e., reinforcement or punishment, are present while the flag is set (Gerstner *et al*., 2018). These additional factors could be implemented by the phasic activity of some neuromodulators, such as dopamine, which is supposed to represent the prediction error (Schultz *et al*., 1997). Dopamine-dependent synaptic plasticity in corticostriatal synapses, which underlies reinforcement learning, shows opposing dopamine dependence for D1 and D2 neurons (Shen et al., 2008; Yagishita *et al*., 2014). Although detailed mechanisms concerning how trace time scales for reinforcement and punishment are modulated remain unclear, our theoretical consideration of the state transition diagram imitating obsessive compulsive symptoms showed that imbalanced trace factors *ν*^+^ > *ν*^−^ could lead to a condition of repetitive choices of abnormal reaction to intrusive thoughts and neutralization of anxiety, similar to obsession and compulsion in OCD (Figure 1a–e). Our experimental data from patients with OCD and HCs strongly support predictions from our computational model, i.e., apparent impairment in learning with delayed feedback (Figure 3b, 3c, and 3d) and extremely imbalanced trace factors *ν*^+^ > *ν*^−^ in OCD (Figure 4a). It is noteworthy that imbalanced trace factors *ν*^+^ > *ν*^−^ are quite convincing because the conventional pathophysiological model of OCD suggests excess tone in the direct pathway over the indirect pathway (Abe *et al*., 2015; Pauls *et al*., 2014; Sakai *et al*., 2011), which are supposed to be related to *ν*^+^ and *ν*^−^, respectively.

Imbalance for positive and negative prediction errors has been often discussed about the learning rate *α*^±^ (e.g. (Lefebvre *et al*., 2017; Palminteri and Lebreton, 2022)). Although we showed that the learning-rate imbalance alone cannot cause OCD-like behavior in the supposed state-transition environments (Figure 1g), it is valuable to clarify the relation between the learning-rate and trace-factor imbalances. The trace-factor imbalance has no effect on contingency of immediate outcomes, while the learning-rate imbalance does affect it (see the theoretical derivation section of STAR Methods). The trace-factor imbalance affects contingency of delayed outcomes through correlation with temporal sequences. In contrast, the learning-rate imbalance affects delayed outcomes uniformly. The learning-rate imbalance can cause risk preference or aversion (Supplementary Figure 2), the mechanisms of which correspond to the discussion by the reference point in prospect theory (Kahneman and Tversky, 1979).

There have been several computational models of OCD, all of which focused primarily on compulsive behaviors (Gillan and Robbins, 2014; Hauser *et al*., 2017; Vaghi *et al*., 2017). Compulsive actions are thought to be reinforced by the rewarding effect of relief from anxiety (Cavedini *et al*., 2006). Excessive anxiety induces excessive reinforcement of compulsion leading to habit formation (Gillan and Robbins, 2014). This excessively habitual compulsion is considered a cause of OCD (Gillan and Robbins, 2014; Gillan and Sahakian, 2015). However, obsessive thoughts that drive anxiety also increase with the severity of OCD symptoms (Cavedini *et al*., 2006). Although the essence of OCD symptoms is the abnormal repetition of obsession and compulsion (Cavedini *et al*., 2006), thus far there has been no unified model to explain growth of both obsession and compulsion. Our unified model successfully represents the vicious circle of obsession and compulsion, which is the phenomenological characteristic of OCD (Cavedini *et al*., 2006) (Figure 1a–1e). Moreover, this model extends our understanding of therapeutic effects of first-line treatments of OCD. While ERP seems to promote appropriate choices to prevent abnormal reactions to intrusive thoughts, even under imbalanced trace factors (Figure 1d), SRIs resolve the imbalance itself (Figure 3d and 4b). In practice guidelines for treatment of OCD (Koran *et al*., 2007), combination therapy of ERP and SRI is recommended if patients do not respond to ERP monotherapy. We identified such a condition in which ERP fails in our computational model (Figure 1f). SRI add-on therapy in this ERP-refractory condition can be viewed as assisting normalization of trace factors. Regarding relationships between the neuromodulator, serotonin, and time scales of the eligibility trace, serotonin seems to modulate synaptic plasticity in many brain regions directly or indirectly through its regulatory effects on other neuromodulatory systems (Brzosko et al., 2019; Cavaccini et al., 2018; Gerstner *et al*., 2018; He *et al*., 2015). In our previous study, we also found similar modulatory effects of trace factor time scales. That is, depletion of the serotonin precursor, tryptophan, increased the *ν*^+^ > *ν*^−^ imbalance, compared with the control condition (Tanaka *et al*., 2009). There is still abundant scope for further research aimed at illuminating how serotonin modulates the time scale of trace factors.

Beyond the range of clinical phenotypes seen in patients with OCD, a broader continuum of the obsessive-compulsive trait is also observed in behavioral characteristics. Specifically, the imbalanced (*ν*^+^ > *ν*^−^) HC cluster showed a significantly greater propensity for Checking and Attention to Detail (Figure 4c). These results increase the reliability of our clinical findings and support the generalizability of our findings to the broader population on the obsessive-compulsive continuum. Further study with a greater focus on patients with OCD and their unaffected healthy first-degree relatives should be performed to evaluate the potential of our findings as an endophenotype of OCD (Chamberlain and Menzies, 2009).

This study provided evidence that one’s own imbalanced implicit learning between reinforcement and punishment can induce a vicious cycle of obsession and compulsion even despite explicit environmental understanding. The discrepancy between explicit environmental knowledge and implicit learning could lead to ego-dystonic anxiety (Gillan and Robbins, 2014). We believe that these results are not limited to OCD, but capture characteristics of an obsessive-compulsive type of anxiety widely present in the general population (Figure 4a and 4c). Anxiety can manifest itself in many different forms. In some situations, anxiety may motivate people to take action or to strive to achieve a goal. In other situations, people may experience various forms of anxiety as a symptom of an anxiety disorder, such as social anxiety disorder or generalized anxiety disorder. Because we cannot cover all forms of anxiety in this research, further study should be performed with a greater focus on constructing computational models of other types of anxiety.

## Conclusions

Although we think that we always make rational decisions, our computational model proves that we sometimes implicitly reinforce maladaptive behaviors and may feel intense anxiety. Psychiatric symptoms are often regarded as mental alterations that are not directly quantifiable, but they can be directly assessed by creating appropriate computational models. Although it is currently difficult to identify treatment-resistant patients based upon their clinical symptoms, our computational model suggests that patients with highly imbalanced trace scale parameters may not respond to behavioral therapy alone. These results suggest that our findings could one day be applied to appropriate selection of OCD treatments. In addition, psychiatric symptoms have been regarded in recent years as a symptom dimension common to various mental diseases rather than to a specific disease. In this study, we focused on patients with OCD and healthy participants, but our approach could be applied to assess the obsessive-compulsive dimension in various populations (Gillan et al., 2019). Future research is needed to address these hypotheses in prospective longitudinal cohorts or in larger cohorts with various psychiatric symptoms.

### Limitations of the study

We did not experimentally test whether the strength of discrepancy between explicit environmental understanding and implicit learning is correlated with the strength of ego-dystonic anxiety. Since this discrepancy is thought to lead to ego-dystonic anxiety (Gillan and Robbins, 2014; Vaghi *et al*., 2017), it was not the focus of this paper. We want to emphasize that our model can explain a spiral of repetitive obsession and compulsion, a core feature of OCD that previous computational models have failed to explain.

We found that participant parameters were densely distributed in the balanced (*ν*^+^ = *ν*^−^) and imbalanced (*ν*^+^ > *ν*^−^) area, which are localized parts of the *ν*^+^ and *ν*^−^ parameter space. There are two possibilities for this. One possibility is that humans tend to exhibit *ν*^+^ = *ν*^−^ and *ν*^+^ > *ν*^−^ parameter settings as universal characteristics. We find that imbalanced HCs show a stronger OCD tendency when compared to balanced HCs (Figure 4c), supporting the possibility that this is a universal characteristic, rather than simple experimental noise. It might be an original human biological limitation that makes us prone to OCD. Furthermore, although we have focused on OCD in this paper as a prominent example of the discrepancy between explicit environmental understanding and implicit learning, the discrepancy is not only associated with OCD, as noted in the Introduction. We believe that minor discrepancies exist in the daily lives of the general population. However, in this paper, we cannot address the reason that imbalanced HCs show a stronger OCD tendency, but do not develop OCD. This needs to be examined in a future study. Another possibility is that this bias might be a tendency in the delayed feedback task. We have confirmed that parameter estimation is accurate in all *ν*^+^ and *ν*^−^ parameter spaces (Supplementary Figure 1), and it is unlikely that such a bias would occur in parameter estimation. However, we cannot completely rule out the possibility of such a bias emerging as a trend in real data. We believe that this issue can be addressed by conducting parameter estimation using a different reinforcement learning task as a future study. Finally, we would like to emphasize that validation using the delayed feedback task is sufficient to find a theoretically derived difference between the OCD and HC groups (Figure 4a).

## Acknowledgments

All authors had full access to data used in this study. The corresponding author, S. C. Tanaka, assumes responsibility for data integrity and accuracy of the data analysis. This study was mainly supported by JSPS KAKENHI Grant Number JP21H05172 (S.C. Tanaka). The following partly supported this study: “Research and development of technology for enhancing functional recovery of elderly and disabled people based on non-invasive brain imaging and robotic assistive devices”, the Commissioned Research of National Institute of Information and Communications Technology (NICT), JAPAN (S.C. Tanaka); JSPS KAKENHI Grant Number JP16K01958 and JP16H06396 (S.C. Tanaka); JP16H01516 and JP18H05524 (Yutaka Sakai); JP16H01512 and the Sakamoto Research Foundation of Psychiatric Diseases (Yuki Sakai); the Joint Usage/Research Center (“Behavioral economics”) of Institute of Social and Economic Research, Osaka University (J. Narumoto).

## Author Contributions

Yuki S, Yutaka S, JN, and SCT designed the study. Yutaka S developed the theory and performed computational modeling. Yuki S, YA, JN, and SCT collected the data. Yuki S conducted the computational modeling and statistical analysis of the data. Yuki S and Yutaka S wrote the manuscript, which all authors edited. SCT supervised the conduct of this study and acquired funding to support theory development and data analysis.

## STAR Methods

## KEY RESOURCES TABLE

| REAGENT or RESOURCE | SOURCE | IDENTIFIER |
| --- | --- | --- |
| Software and algorithms |  |  |
| Python | Python | https://www.python.org/ |
| MATLAB | MathWorks | https://www.mathworks.com/ |
| R | R Development Core Team | https://www.r-project.org/ |
| Original code (theoretical derivation of conditions for OCD-like behavior) | This paper | The theoretical derivation section of STAR Methods |
| Original code (parameter estimation) | This paper | [https://zenodo.org/record/6879513#.](https://zenodo.org/record/6879513#.YtoPcITP02-)<br>YtoPcITP02-<br>(DOI:<br>10.5281/zenodo.6879513) |

## RESOURCE AVAILABILITY

### Lead contact

Further information should be directed to and will be fulfilled by the lead contact, Saori C. Tanaka.

### Materials availability

Materials are available from the corresponding author upon reasonable request.

### Data and code availability

- De-identified human behavioral data of participants giving consent to public data release except for patients with OCD will be shared by the lead contact upon reasonable request. Patient data supporting conclusions of this paper are not publicly available because they contain information that could compromise research participant privacy or consent.
- All original code will be publicly available as of the date of publication (https://zenodo.org/record/6879513#.YtoPcITP02– (DOI: 10.5281/zenodo.6879513) or https://github.com/YukiSakai1209/Sakai-and-Sakai-et-al._CellReports_2022).
- Any additional information required to reanalyze the data reported in this paper is available from the lead contact upon request.

## EXPERIMENTAL MODEL AND SUBJECT DETAILS

### Participants

Forty-five patients with OCD and 168 HCs participated in the behavioral task. The Medical Committee on Human Studies at Kyoto Prefectural University of Medicine (KPUM) and the Ethics Committee at Advanced Telecommunications Research Institute International (ATR) approved all procedures in this study. All participants gave written informed consent after receiving a complete description of the study. All methods were carried out following approved guidelines and regulations. Trained, experienced clinical psychiatrists and psychologists assessed all participants.

All patients were primarily diagnosed using the Structured Clinical Interview for DSM-IV Axis I Disorders-Patient Edition (SCID) (First et al., 1994). Experienced clinical psychiatrists or psychologists applied the Yale-Brown Obsessive-Compulsive Scale (Y-BOCS) (Nakajima et al., 1995), the 17-item Hamilton Depression Rating Scale (HDRS) (Hamilton, 1967), and Wechsler Adult Intelligence Scale - Third Edition (WAIS-III) for evaluation of obsessive-compulsive symptoms, depressive symptoms, and IQ in patients with OCD, respectively. There was one missing value for the HDRS. Exclusion criteria were 1) a cardiac pacemaker or other metallic implants or artifacts; 2) significant disease, including neurological diseases, disorders of the pulmonary, cardiac, renal, hepatic, or endocrine systems, or metabolic disorders; 3) prior psychosurgery; 4) IQ < 80 using WAIS-III; 5) DSM-IV diagnosis of mental retardation or pervasive developmental disorders based on a clinical interview and psychosocial history, and 6) pregnancy. We excluded 3 OCDs due to IQ < 80 using the WAIS-III. Seventeen HCs were excluded for medical or experimental reasons. Subsequent analysis was conducted in 42 patients with OCD and 151 HCs. There were no significant differences in age, sex, or handedness (Supplementary Table 1). To the extent possible, we selected patients without a current DSM-IV Axis I diagnosis of any significant psychiatric illness except OCD. Only two patients with trichotillomania, one patient with a tic disorder, two patients with panic disorders, and one patient with bulimia nervosa were included as patients with comorbidities. Handedness was classified based on a modified 25-item version of the Edinburgh Handedness Inventory. Thirteen patients with OCD were drug-free for all types of psychotropic medication. SRI and imipramine equivalent doses (Inada and Inagaki, 2015) of the remaining 29 patients are summarized in Supplementary Table 2.

## METHOD DETAILS

### Separate eligibility trace model

Eligibility traces determine the weight assignment of outcome prediction errors for recent actions (Figure 1b). If eligibility traces for positive and negative prediction errors are implemented in distinct neural systems, the respective eligibility traces 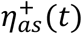 and 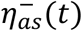 at time step *t* for action *a* (*a* = 0, 1, …) in state *s* (*s* = 0, 1, …) obey the following equation (Sutton and Barto, 1998),

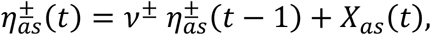

where *X*_*as*_ (*t*) = 1 when action *a* is chosen in state *s* at time step *t* and *X*_*as*_ (*t*) = 0 otherwise. The factors *ν*^±^ (0 < *ν*^±^ < 1) determine the decaying time scales of the eligibility traces. The policy parameters *q*_*as*_(*t*) (*a* = 0, 1, …) determine choice probability in state *s* as a soft-max function,

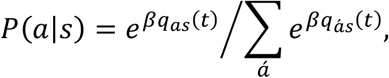

where the greediness parameter β represents the degree of the exploration-exploitation balance. Each policy parameter is updated as below,

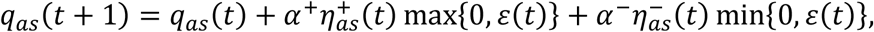

where *ε*(*t*) denotes the outcome prediction error and max{0, *ε*(*t*)} and min{0, *ε*(*t*)} represent positive and negative components of the prediction error, respectively (Figure 1b). Theoretically, the trace factors *ν*^±^ and the learning rates *α*^±^ should be balanced as *ν*^+^ = *ν*^−^ and *α*^+^ = *α*^−^.

However, separate neural implementation makes it difficult to maintain the balance perfectly. Basically, we assumed that the balance of trace factors could be broken: *ν*^+^≠ *ν*^−^, under the balanced learning rates *α*^+^ = *α*^−^. Only for the control comparison, we considered the learning-rate imbalance.

The outcome prediction error *ε*(*t*) is determined as a function of the current outcome *r*(*t*) and the policy parameters {*q*_*as*_}. The form depends on learning algorithms. In actor-critic learning, the outcome prediction is based on the state value *ν*_*s*_ = ∑_*a*_ *q*_*as*_, and the prediction error *ε*(*t*) = *r*(*t*) + *γν*_*s*(*t*+1)_ – *ν*_*s*(*t*)_, where *r*(*t*) denotes the current outcome and *s*(*t*) and *s*(*t* + 1) denote the current and next states. The parameter *γ* denotes the discount factor, which determines the weight of future prediction. In SARSA and Q-learning, the outcome prediction is based on the action value estimated as each policy parameter *q*_*as*_, and the prediction error *ε*(*t*) = *r*(*t*) + *γV*_*s*(*t*+1)_ – *q*_*s*(*t*)*s*(*t*)_. The difference is in the term of the next state value: *V*_*s*(*t*+1)_ = *q*_*s*(*t*+1)*s*(*t*+1)_ in SARSA and 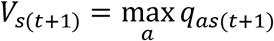 in Q-learning.

### State transition diagram imitating OC symptoms

We described mental states involved in obsessive-compulsive symptoms in stochastic transitions between two states: a relief state (*s*=0) and an obsessive state (*s*=1) (Figure 1a). We assumed two options in each state: “compulsion” (*a*=1) and “other” (*a*=0) in the obsessive state, and “abnormal reaction to intrusive thoughts” (*a*=1) and “other” (*a*=0) in the relief state. We defined a matrix *b* to determine the state-transition probabilities,

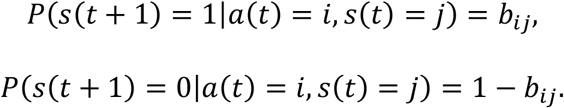

We assumed that every stay in the obsessive state produced a negative outcome normalized to −1. The relative cost of compulsion was denoted as *c*. Although *b*_00_>0 allows passive anxiety, we focused on *b*_00_=0 in the main article. More general cases, including passive anxiety, are considered in the theoretical derivation section of STAR Methods.

### Demonstration of OCD-like behavior

For the example in Figure 1, we fixed the parameters of the state transition as *b*_00_=0, *b*_10_=1, *b*_01_=0.9, *b*_1_=0.5, and *c*=0.01 and the learning parameters as α=0.1, *β*=1, *γ* = 0.5, and *ν*^+^=0.8, except for *ν*^−^. For Figure 1c, the trace factor for punishment was set as *ν*^−^=0.6 and the initial values of variables as 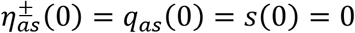. The simulation consisted of 100,000-time steps. For Figure 1d, we set the trace factors as *ν*^−^=0.1 and the initial values as 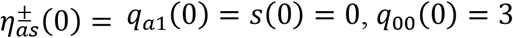, and *q*_10_(0) = −3, to show that reinforcement of the abnormal reaction to intrusive thoughts started from even a low reaction rate. After 50,000-time steps, compulsion (*a*=1 in *s*=1) was always prevented, and other (*a*=0 in *s*=1) was forced regardless of the choice probability, demonstrating the behavioral therapy of ERP.

For Figure 1e and f, we evaluated the fraction of 100 simulation runs in which the reaction rate to intrusive thoughts was reinforced from a low reaction rate on 40 × 40 grids in (*ν*^+^, *ν*^−^) space. In each simulation, 200 instances of forced abnormal reaction were intermittently caused in the relief state because spontaneous abnormal reactions scarcely occurred at a low reaction rate. The simulation was judged to be abnormal-reaction-reinforced if the choice probability of reaction to intrusive thoughts became larger than the initial value. The initial values of variables were set as 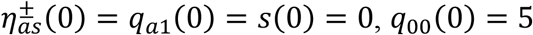, and *q*_10_(0) = −5. Each forced abnormal reaction was caused when 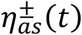 sufficiently approached zero in the relief state 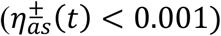.

For Figure 1g and h, we evaluated the fraction of abnormal-reaction reinforcements as in Figure 1e and f, but the learning-rate imbalance *α*^+^/(*α*^+^ + *α*^−^) and the value of the greediness parameter *β* were swept for the balanced trace factors *ν*^+^=*ν*^−^=0.8. The scale of the learning rate is independent of the steady behavior, and excessively large learning rate will practically cause instability (Sutton and Barto, 1998). Therefore, we swept only the ratio of the learning rates under the fixed summation *α*^+^ + *α*^−^=0.1. Grids for *β* are arranged in logarithmic scale in the range from 10^0^=1 to 10^2^=100.

### Behavioral task

The delayed feedback task was similar to our previous research (Tanaka *et al*., 2009), except for the presented stimuli. Participants chose one of the two options (abstract cues) displayed on the screen by pressing a left or right button in each trial within 1 s after an auditory cue (Figure 2a). Depending on the selected stimulus, monetary feedback with different outcomes (+10, +40, −10, or −40 yen) was displayed either immediately after the button press or three trials later (Figure 2b and c). We did not offer monetary feedback for the first five trials because participants can learn the relationships between stimuli and outcomes or delays quickly. If participants pressed a button before the auditory cue or more than 1 s passed without any button press, −50 yen was displayed as a punishment. Such trials were considered error trials, and delayed outcomes were not considered.

Two abstract cues were displayed at each trial side by side on the screen (Figure 2a). We prepared 16 pairs (Figure 2b), counterbalancing the number of appearances of each stimulus. The 16 pairs of stimuli were presented in pseudorandom order. Each pair was presented as the scheduled number of trials in each session: each of six pairs [+10(0) vs. +40(0); +10(3) vs. +40(3); −10(0) vs. −40(0); −10(3) vs. −40(3); +10(0) vs. +40(3); −10(0) vs. −40(3)] was presented in 10 trials during a single session, and each of 10 pairs [+40(0) vs. +40(3); +10(0) vs. +10(3); −10(0) vs. −10(3); −40(0) vs. −40(3); +10(3) vs. +40(0); −10(3) vs. −40(0); +10(0) vs. −10(0); +40(0) vs. −40(0); +10(3) vs. −10(3); +40(3) vs. −40(3)] was presented in five trials during a single session. Each participant performed 110 trials during a single session and six sessions in each experiment. About 28 min was required for participants to complete six sessions. At the beginning of each session, the session number was displayed on the screen for 2.5 s. Before the task, each participant practiced the test session under the same task settings except for stimuli, and we confirmed that all participants understood the task set.

## QUANTIFICATION AND STATISTICAL ANALYSIS

### Statistical analysis of the behavioral task

The total outcome (except for punishments related to button press errors), reaction time, and the number of error trials were compared between the OCD and HC groups. Patients with OCD showed a significantly lower total monetary outcome than HCs [median (interquartile range): patients with OCD, 2705 (1510–3542) yen; HCs, 4700(3095–5955) yen; Brunner-Munzel test, statistic = 5.31, p < 0.001], whereas there were no group differences in reaction time [median (interquartile range): patients with OCD, 545.9 (503.4–614.8) ms; HCs, 522.6 (475.9– 574.8) ms; Brunner-Munzel test, p > 0.05)] and numbers of button press errors [median (interquartile range): patients with OCD, 4.5 (2.25–14.75); HC, 4.0 (2.0–8.0); Brunner-Munzel test, p > 0.05). These results suggested impaired learning in individuals with OCD compared with HCs.

Based on our hypothesis of trace factors *ν*^+^/*ν*^−^, patients with OCD were expected to show impaired learning in stimuli with delayed feedback. To confirm relationships between the parameters (*ν*^+^ and *ν*^−^) and the total outcome from stimuli with immediate or delayed feedback, we conducted the simulation. We evaluated the total outcome from stimuli with immediate or delayed feedback using actor-critic learning with separate eligibility traces in the same experimental settings as the actual experiment. We created 500 simulated cases using each combination of *ν*^+^ and *ν*^−^ (0-0.99 in 0.01 increments). Values of *α* and *β* were fixed as the median values of HC estimated parameters (α, 0.0011; *β*, 2.96). We projected the mean values of total simulated outcome from stimuli with immediate or delayed feedback in *ν*^+^/*ν*^−^ space (Figure 3a).

An ANOVA-type statistic (ATS) for nonparametric repeated measures ANOVA (Noguchi *et al*., 2012) was conducted to clarify between-group differences. Bonferroni-Holm–corrected post-hoc comparisons using Brunner-Munzel test were conducted when needed. Therapeutic effects of SRIs were evaluated using Spearman’s rank correlation between the dose equivalence of SRIs (Inada and Inagaki, 2015) (Supplementary Table 2) and the total outcome from stimuli with immediate or delayed feedback in sessions 5 and 6.

### Model validation and parameter estimation

We fitted behavioral data with actor-critic learning with separate eligibility traces for positive and negative prediction errors. We defined 16 states for possible pairs of stimuli presented at each trial. Available actions at each state involved choosing alternative stimuli. To facilitate model-fitting in the face of limited experimental data from each participant, we used regularizing priors that favored realistic values and maximum a posteriori estimation rather than maximum likelihood estimation (Daw, 2011). The learning rate *α* and trace factor *ν* were constrained to the range of 0 ≤ *α* ≤ 0.95 and 0 ≤ *ν* ≤ 0.95 with a uniform prior. The exploration-exploitation degree *β* was constrained to the range of 0 ≤ *β* ≤ 100 with a gamma (2,3) prior distribution that favored relatively lower values. We fixed the discount factor *γ* = 0, because the term of the next state value was just noise in our delayed feedback task in which the state of each trial was randomly selected. We optimized parameters by minimizing the negative log posterior of the data with different parameter settings using the hyperopt package (Bergstra et al., 2011).

Likelihood ratio tests to assess the contribution of the additional parameter in our model (four parameters: *α, β, ν*^+^, *ν*^−^) compared with the standard actor-critic learning model (three parameters: *α, β, ν*) showed that the additional parameter was justified in 133 of 193 participants (*X*^2^ test with one degree of freedom, p < 0.05). In addition, we compared the separate eligibility trace model (four parameters: *α, β, ν*^+^, *ν*^−^) and the separate learning rate model (four parameters: *α*^+^, *α*^−^, *β, ν*). As these models have the same number of parameters, we compared their posterior likelihoods directly. This showed that the separate eligibility trace model provided a better fit than the separate learning rate model in 95.3% of all participants (184/193). To further validate the actor-critic learning model with separate eligibility traces, we simulated the behavioral data of each group using the estimated parameters. We randomly chose 100 participants from each group with duplicates and simulated the total outcome using estimated parameters of the selected participants. We illustrated the mean and standard errors of the simulated total ouctome (Figure 3e, f) and compared them with the experimental results (Figure 3b and 3c). We confirmed that the specific impairment in learning from stimuli with delayed feedback in the simulated OCD group (Figure 3e and 3f) is similar to the experimental data (Figure 3b and 3c). Because our target model was validated with behavioral data, we applied the separate eligibility trace model in the substantive analysis.

We evaluated the performance of parameter estimation in actor-critic learning with separate eligibility traces. We created 200 simulation data points using a model with the following parameters: *α*, 0.1 ± 0.05 (mean ± SD); *β*, 1 ± 0.2; *ν*^+^ and *ν*^−^, randomly selected within 0.01–0.95. Parameter estimation was quite accurate (Yazdani et al., 2018) (Supplementary Figure 1). Specifically, Pearson’s r and the mean absolute error between the true and estimated *ν*^+^ or *ν*^−^ were 0.99 and 0.03, respectively (Supplementary Figure 1).

We compared the *ν*^+^/*ν*^−^ distribution in our model between groups using PERMANOVA with the ADNOIS function and 10,000 permutations using the Euclidean distance implemented in the statistical package R (Anderson et al., 2017). Multivariate homogeneity of group dispersions was confirmed with the *betadisper* function with 10,000 permutations in R (Anderson *et al*., 2017).

The learning rate *α* and inverse temperature *β* were compared using the Brunner-Munzel test. To further investigate relationships between clinical characteristics and estimated parameters in the HC group, we conducted cluster analysis using HDBSCAN (Campello *et al*., 2015). We detected two clusters (balanced cluster, n = 83; imbalanced cluster, n = 57; the remaining 11 HCs were not clustered) and evaluated their obsessive-compulsive trait using the five PI subscales: “Checking”, “Dirt”, “Doubt”, “Impulse”, “Precision” (Sugiura and Tanno, 2000). In addition, the propensity to adhere to fine-grained details was evaluated using the Attention to Detail subscale of AQ (Wakabayashi *et al*., 2004). There were 14 missing values in PI (n = 126) and 1 missing value in AQ (n = 140). Therapeutic effects of SRIs were evaluated using Spearman’s rank correlation between the SRI dose and imbalanced settings of *ν* (*ν*^+^ − *ν*^−^).

### Theoretical derivation of OCD-like behavior

Here, we analytically derived conditions in which the separate eligibility trace model (Figure 1b in the main text) exhibits OCD-like behavior in the obsession-relief state transition (Figure 1a in the main text). In the following derivations, our manual calculations were confirmed using the MATLAB Symbolic Math Toolbox (Mathworks, version R2017a). Hence, we did not include the derivation passages in the manuscript, because they are too long and complicated. Instead, we provide the MATLAB source code to confirm the equations (see the section of MATLAB script to confirm derivations).

#### 1. Stationary analysis of the state transition

The state transition diagram imitating obsessive compulsive symptoms (Figure 1a) consists of a relief state (*s*_0_ = 0) and an obsessive state (*s*_1_ = 1). Available options are “abnormal reaction to intrusive thoughts” (*a*_1_ = 1) and “other” (*a*_0_ = 0) in *s*_0_ and “compulsion” (*a*_1_ = 1) and “other” (*a*_0_ = 0) in *s*_1_. Define a binary vector variable *x*(*t*) to represent the combination of state and action, (*s*(*t*),*a*(*t*)), at time step *t* as

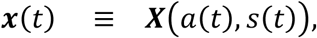

where *X*(*a, s*) is a binary vector function defined as

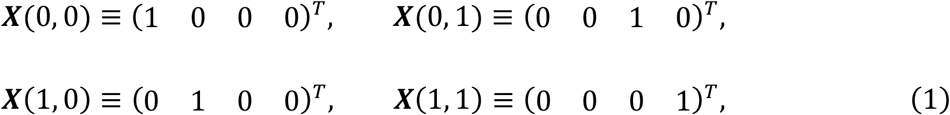

where *M*^*T*^ means transposition of the matrix *M*. The expectation value of vector *x, E*[*x*] represents the probability distribution of the combination of state and action. Throughout the theoretical derivation section of STAR Methods, all four-dimensional vectors are denoted as column vectors, each of which consisted of components corresponding to state-action patterns in the order *a*_0_*s*_0_, *a*_1_*s*_0_, *a*_0_*s*_1_, and *a*_1_*s*_1_.

Likewise, let *s*(*t*) be a binary vector variable to represent only state:

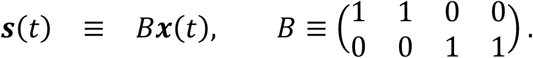

The combination of state and action at time *t* determines the state transition probabilities,

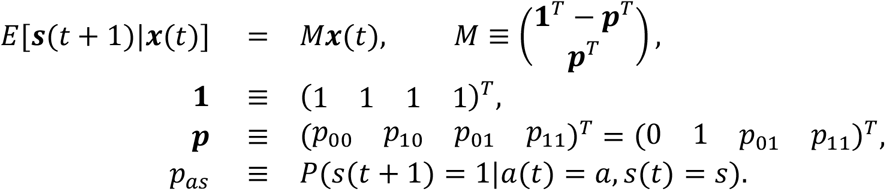

Let 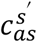 be a given cost at a state transition: *a*(*t*) = *a* and *s*(*t*) = *s* to *s*(*t* + 1) = *s*^′^:

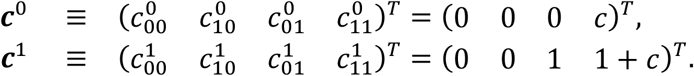

We focused on the case in which a compulsive action in the obsessive state minimizes immediate cost. The expected cost for the compulsion is 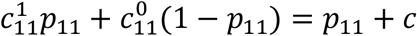, and the expected cost for “other” is 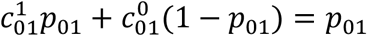. Thus, we focused on the condition:

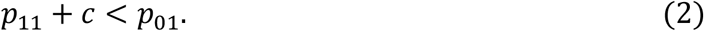

The conditional expectation value of *x*(*t*) on state *s*(*t*) is described with the state-dependent choice probabilities,

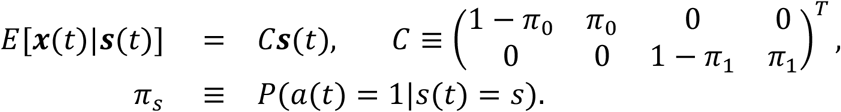

Markov chains for variables *s* and *x* are respectively described as

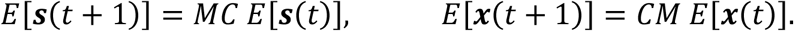

The stationary distribution of *s* is obtained as the eigenvector of 2 × 2 matrix *MC* satisfying the recursive equation *E*[*s*] = *MC E*[*s*]. The stationary distribution of *x* is obtained as *E*[*x*] = *C E*[*s*]. The average outcome is described as

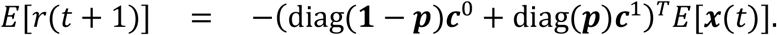

The partial derivative for *π*_0_ is obtained as (see the section of MATLAB script to confirm derivations)

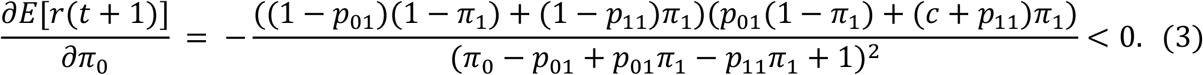

Thus, the optimal reaction rate to intrusive thoughts is always *π*_0_ = 0. The partial derivative for *π*_1_ is obtained as (see the section of MATLAB script to confirm derivations)

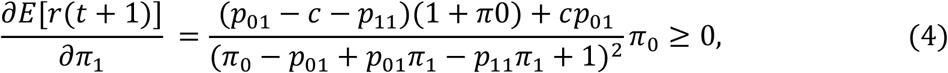

because we focused on the condition (2). Thus, the optimal compulsion rate is *π*_1_ = 1.

#### 2. Stationary analysis of the candidate model

In the separate eligibility trace model, the trace patterns are described as a decaying summation of the past *x* (see the **Separate eligibility trace model** section of STAR Methods).

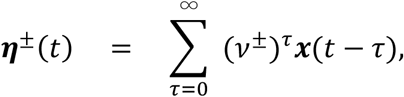

where the trace factors are in the range 0 ≤ *ν*^±^ < 1. We define a policy vector ***q*** to represent the policy parameters,

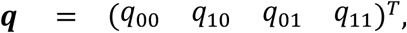

and the conditional expectation value of the policy update Δ***q***(*t*) on *x*(*t*) is described as

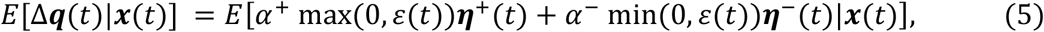

where the outcome prediction error *ε*(*t*) depends on learning algorithms,

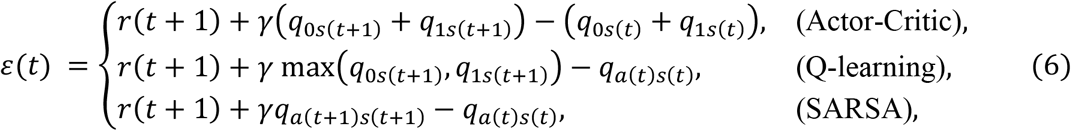

where *r*(*t*) is the outcome at time step *t*. The parameter *γ* denotes the discount factor (0 ≤ *γ* < 1) in the temporal difference (TD) learning framework. The average prediction error should approach zero at the stationary phase after sufficient learning. Therefore, the transition to the relief state *s*_0_ produces a positive error, and the transition to the obsessive state *s*_1_ produces a negative error. Accordingly, the conditional expectation values of the positive and negative prediction errors are described as

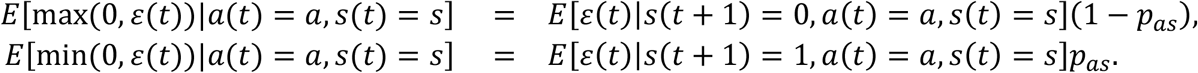

Define vector 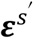 as

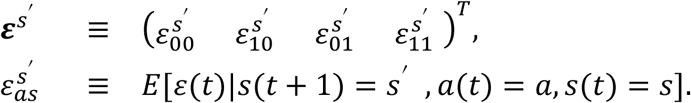

Using these vector forms, the expectation value of the policy update *E*[Δ***q***] is obtained by averaging Eq. (5) over the stationary distribution *E*[*x*(*t*)] = *E*[*x*] as

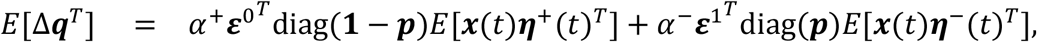

where matrices *E*[*x*(*t*)***η***^±^(*t*)^*T*^] are calculated as

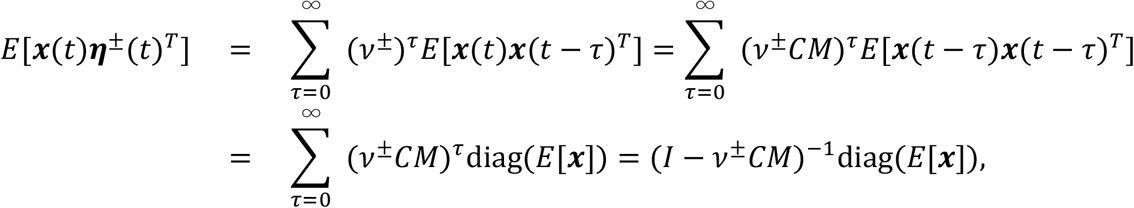

because vector *x*(*t* − *τ*) consists of exclusive binary components. The matrix *I* denotes the identity matrix. Thus, we obtain the expectation value of the policy update as

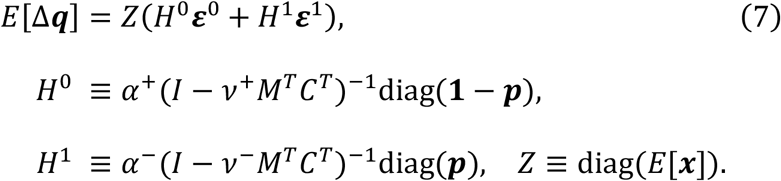

#### 3. Stability of optimal behavior: Q-learning and SARSA

Abnormal-reaction and compulsion rates, *π*_0_ and *π*_1_, are described as soft-max functions of the policy parameters (see the **Separate eligibility trace model** section of STAR Methods),

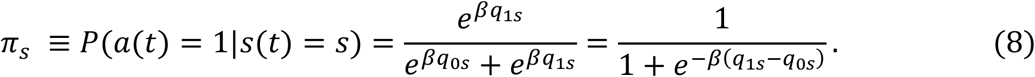

In Q-learning and SARSA, the policy parameters represent action values. The balance of exploration and exploitation is explicitly controlled, and the parameter *β* increased to the limit *β* → ∞ in order to reach the final behavior. Therefore, the abnormal reaction emerges as final behavior rather than exploratory behavior when the difference of the action values in the relief state are positive, *q*_10_ − *q*_00_ > 0. We derive the condition of this inequality around the optimal behavior (*π*_0_, *π*_1_) = (0, 1). The prediction error patterns in Q-learning around the optimal behavior are described as 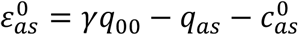, and 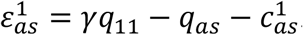. These can be described in vector forms by using the binary vector function *X*(*a, s*) in Eq.(1), the unit vector **1**, and the identity matrix *I*,

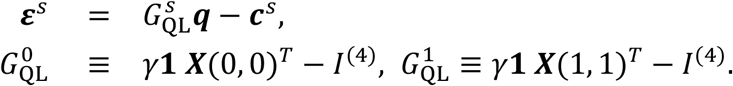

In SARSA, the prediction error patterns are described as

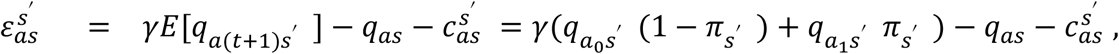

which can be described in vector forms,

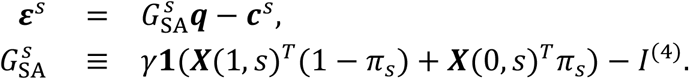

The stationary solution of *E*[Δ***q***] = **0** in Eq. (7) is obtained as

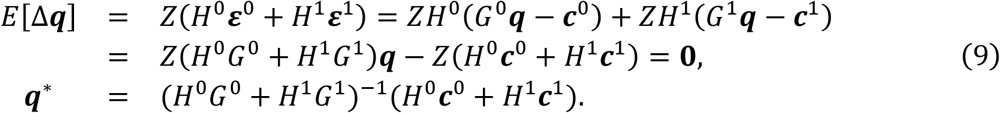

For both Q-learning and SARSA, the difference of the action values in the relief state around the optimal behavior (*π*_0_, *π*_1_) = (0, 1) are obtained as (see the section of MATLAB script to confirm derivations)

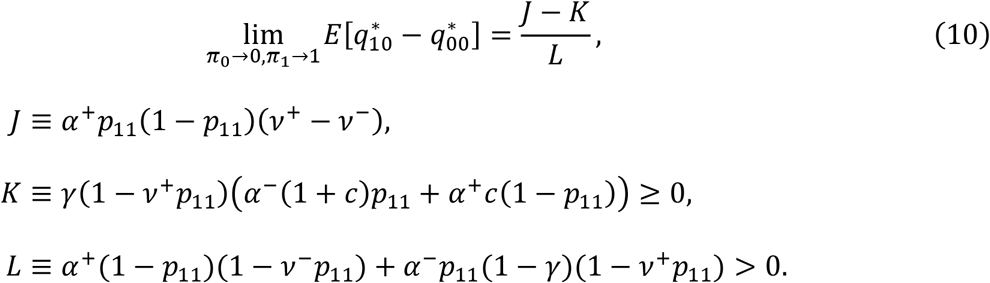

If *J* > *K*, then the zero abnormal-reaction rate to intrusive thoughts *π*_0_ = 0 is broken. Thus, the condition in which the OCD-like behavior might emerge is described as

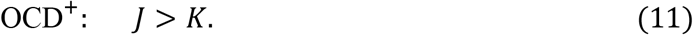

Because *K* ≥ 0, a necessary condition is that *J* > 0, namely, a trace imbalance: *ν*^+^ > *ν*^−^.

In behavioral therapy [exposure and response prevention (ERP)], compulsion is prevented in the obsessive state, that is, *π*_1_ = 0. For Q-learning, the action value *q*_1_ can be used even if the compulsive action (*a* = 1) is prevented in the obsessive state (*s* = 1) by the definition of the algorithm. However, response prevention is equivalent to the infinite response cost, which leads to inequality in action values, *q*_1_ < *q*_01_. Thus, the matrix 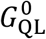 is replaced in the behavioral therapy,

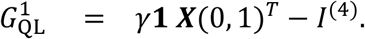

For both Q-learning and SARSA, the difference in action values at the optimal behavior in the behavioral therapy, (*π*_0_, *π*_1_) = (0,0), is obtained as (see the section of MATLAB script to confirm derivations)

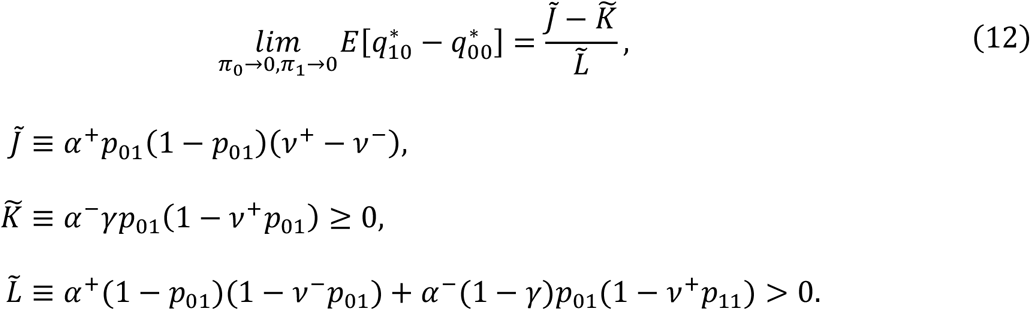

Using the same logic as the derivation of the condition around (*π*_0_, *π*_1_) = (0, 1), the condition in which the behavioral therapy (ERP) might be ineffective is

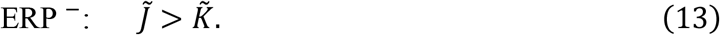

#### 4. Stability of optimal behavior: Actor-critic

In actor-critic learning, policy parameters diverge to approach an optimal deterministic behavior, whereas the state values ***ν*** = *B****q*** converge. The prediction error patterns in actor-critic learning are described as

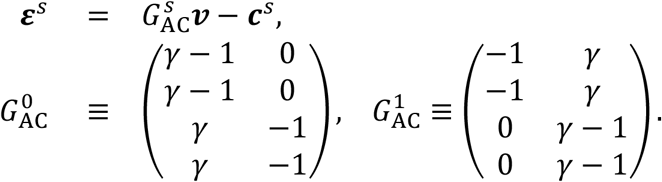

The solution of *E*[Δ***ν***] = *E*[Δ*B****q***] = **0** in Eq. (7) is obtained as

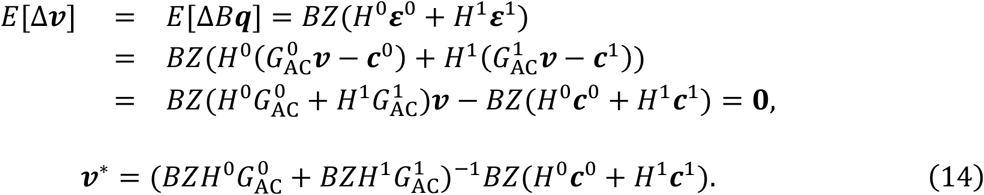

The signs of updates in the abnormal-reaction and compulsion rates for the stationary ***ν***^∗^ determine the stability of the optimal behavior (*π*_0_, *π*_1_) = (0, 1); hence, signs of updates in the abnormal-reaction and compulsion rates coincide with signs of the difference between policy updates Δ*q*_1*s*_ − Δ*q*_0*s*_. To examine learning stability, the difference update is evaluated,

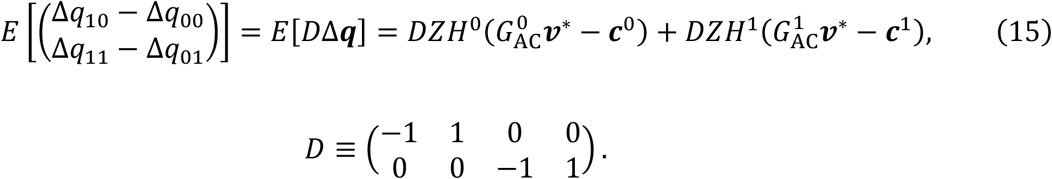

The difference update *E*[*D*Δ***q***] is zero at the optimal behavior (*π*_0_, *π*_1_) = (0, 1) (see the section of MATLAB script to confirm derivations). Therefore, we derived the Taylor expansion up to the first non-zero order (see the section of MATLAB script to confirm derivations),

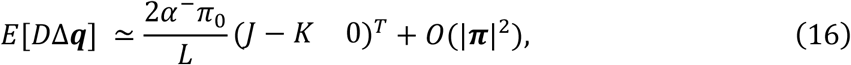

For a sufficiently small abnormal-reaction rate to intrusive thoughts *π*_0_ ≃ 0, the update of the abnormal-reaction rate *0*(*π*_0_) is dominant over the compulsion update *0*(|***π***|^2^). Therefore, the stability of the optimal behavior (*π*_0_, *π*_1_) = (0, 1) would be broken if *E*[Δ*q*_10_ − Δ*q*_00_] > 0. Thus, the condition for stability broken in actor-critic learning is equivalent to those in Q-learning and SARSA, *J* > *K*, as Eq.(11).

In behavioral therapy (ERP) where *π*_1_ = 0 is forced, the difference update *E*[*D*Δ***q***] is also zero at *π*_0_ = 0 (see the section of MATLAB script to confirm derivations). Therefore, we derived the Taylor expansion up to the first non-zero order around *π*_0_ = 0 (see the section of MATLAB script to confirm derivations),

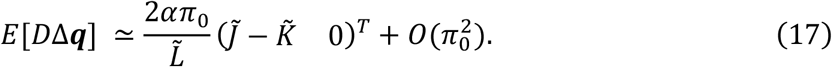

Thus, the condition in which behavioral therapy (ERP) might be ineffective in actor-critic learning is equivalent to that in Q-learning and SARSA, 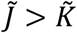, as Eq.(13).

#### 5. Necessary condition for patients with OCD

The state transition diagram (Figure 1a in the main text) represents a certain obsessive-compulsive situation. State-transition probabilities *p*_01_, *p*_11_ and the relative cost of compulsive action *c* depend on the concrete contents of the abnormal reaction to intrusive thoughts, obsessive state, and compulsive action. Patients with OCD exhibit various symptoms in response to many possible situations. Hence, it is significant to derive the condition for the learning parameters in which some state-transition probabilities *p*_01_, *p*_11_ and cost *c* exist to satisfy the condition for OCD-like behavior (11).

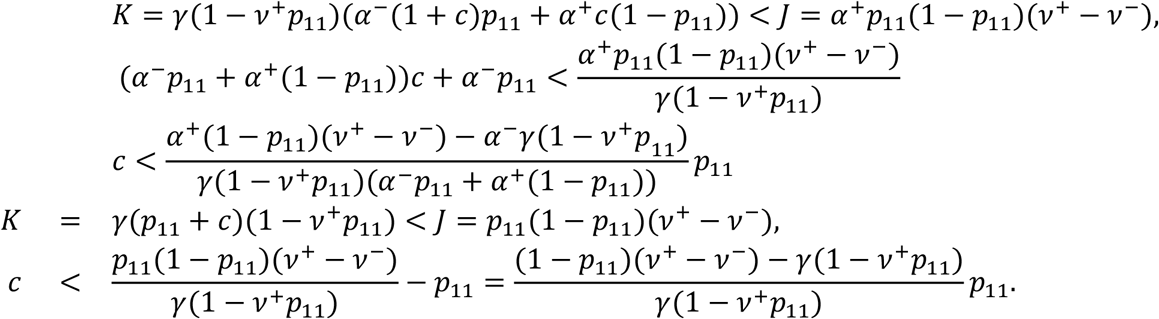

Therefore, the condition for the existence of *c* > 0 is

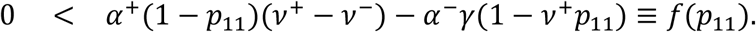

The right-hand value at *p*_11_ = 1 is negative, *f*(1) = −*α*^−^*γ*(1 − *ν*^+^) < 0; hence, the condition for the existence of *p*_11_, such that 0 < *p*_11_ < 1 and *f*(*p*_11_) > 0 is *f*(0) = *α*^+^(*ν*^+^ − *ν*^−^) − *α*^−^*γ* > 0, that is,

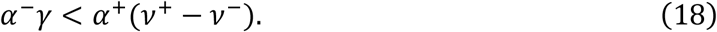

If an individual’s learning parameters satisfy the condition (18), the individual might exhibit OCD symptoms in some situations.

#### 6. MATLAB script to confirm derivations

Copy and execute the following MATLAB script, and you can confirm the equations above mentioned. This script requires Symbolic Math Toolbox.

~~~
% derivation.m
%
% MATLAB script to confirm derivations in Supplementary Note of the article (Sakai & Sakai et al.)
% This requires Symbolic Math Toolbox
%
%%%%% Definitions
syms p01 p11 pi0 pi1 gam np nm ap am al c;
B=[1 1 0 0;0 0 1 1]; p=[0 1 p01 p11]; M=[1-p;p]; C=[1-pi0 0;pi0 0;0 1-pi1;0 pi1];
D=[-1 1 0 0;0 0 −1 1]; c0=[0;0;0;c]; c1=[0;0;1;1+c];
%%%%% Stationary probability distribution
MC=M*C; s=[MC(1,2);1-MC(1,1)]; s=s/sum(s); x=C*s; Z=diag(x);
H0=ap*inv(eye(4)-np*M.’*C.’)*diag(1-p); H1=am*inv(eye(4)-nm*M.’*C.’)*diag(p);
%%%%%%% Optimal Behavior r=simplify(-(diag(1-p)*c0+diag(p)*c1).’*x) %%
Partial derivative of E[r] for pi0 dr0=-((1-p01)*(1-pi1)+(1-p11)*pi1)*(p01*(1-
pi1)+(c+p11)*pi1)/(pi0-p01+p01*pi1-p11*pi1+1)^2;
%% Partial derivative of E[r] for pi1 dr1=pi0*((p01-c-
p11)*(1+pi0) +c*p01)/(pi0-p01+p01*pi1-p11*pi1+1)^2;
checkEq3=simplify(dr0-diff(r,pi0))
checkEq4=simplify(dr1-diff(r,pi1))
%%%%%%% Stationary analysis of SARSA
Gsa0=gam*ones(4,1)*[1-pi0 pi0 0 0]-eye(4); Gsa1=gam*ones(4,1)*[0 0 1-pi1 pi1]-eye(4);
F=H0*Gsa0+H1*Gsa1; qstar=simplify(inv(subs(F,[pi0 pi1],[0 1]))*subs(H0*c0+H1*c1,[pi0 pi1],[0 1]));
J=ap*p11*(1-p11)*(np-nm); K=gam*(1-np*p11)*(am*(1+c)*p11 + ap*c*(1-p11));L=ap*(1-p11)*(1-
nm*p11)+am*p11*(1-gam)*(1-np*p11); checkEq10=simplify(qstar(2)-qstar(1)-(J-K)/L)
%%%%% Behaviral therapy of SARSA qstar=simplify(inv(subs(F,[pi0 pi1],[0
0]))*subs(H0*c0+H1*c1,[pi0 pi1],[0 0])); tJ=ap*p01*(1-p01)*(np-nm); tK=am*gam*p01*(1-
np*p01); tL=ap*(1-p01)*(1-nm*p01)+am*p01*(1-gam)*(1-np*p01);
checkEq12=simplify(qstar(2)-qstar(1)-(tJ-tK)/tL)
%%%%%%% Stationary analysis of Q-learning
Gql0=gam*ones(4,1)*[1 0 0 0]-eye(4); Gql1=gam*ones(4,1)*[0 0 0 1]-eye(4); F=H0*Gql0+H1*Gql1;
qstar=simplify(inv(subs(F,[pi0 pi1],[0 1]))*subs(H0*c0+H1*c1,[pi0 pi1],[0 1])); checkEq10=simplify(qstar(2)-
qstar(1)-(J-K)/L)
%%%%% Behaviral therapy of Q-learning
Gql1=gam*ones(4,1)*[0 0 1 0] - eye(4); F=H0*Gql0+H1*Gql1;
qstar=simplify(inv(subs(F,[pi0 pi1],[0 0]))*subs(H0*c0+H1*c1,[pi0 pi1],[0
0])); checkEq12=simplify(qstar(2)-qstar(1)-(tJ-tK)/tL)
%%%%%%% Stationary analysis of Actor-Critic
Gac0=[gam-1 0;gam-1 0;gam −1;gam −1]; Gac1=[−1 gam;-1 gam;0 gam-1;0 gam-
1]; v=simplify(inv(B*Z*(H0*Gac0+H1*Gac1))*B*Z*(H0*c0+H1*c1)); %%%
Stationary state value
Ddq=simplify(D*Z*(H0*(Gac0*v-c0)+H1*(Gac1*v-c1))); %%% Policy update difference for stationary state
value %%%%% Tailor expansion of E[D Delta q]
Ddq_opt=simplify(subs(Ddq,[pi0 pi1],[0 1])) %%% Policy update differences at pi=[0,1] are 0
%%% Partial derivertives of policy update difference at pi=[0,1]
Ddq_0=simplify(subs(diff(Ddq,pi0),[pi0 pi1],[0 1])); Ddq_1=simplify(subs(diff(Ddq,pi1),[pi0 pi1],[0 1]))
checkEq16=simplify(Ddq_0-[2*am*(J-K)/L; 0])
%%%%% Behaviral therapy of Actor-Critic
Ddq_beh=subs(Ddq,pi1,0);
%%%%% Tailor expansion of E[D Delta q]
Ddq_opt=simplify(subs(Ddq_beh,pi0,0)) %%% Policy update differences at pi0=0 are 0
%%% Derivertives of policy update difference at pi0=0
Ddq_0=simplify(subs(diff(Ddq_beh,pi0),pi0,0)); checkEq17=simplify(Ddq_0-
[2*am*(tJ-tK)/tL; 0])
~~~

#### 7. Learning-rate and trace-factor imbalances

In this study, we supposed the imbalance in the downstream of the common outcome prediction error *ε*(*t*). The downstream was assumed to be separated for the sign of *ε*(*t*). We introduced the separate eligibility trace model, in which each policy parameter is updated below (see the **Separate eligibility trace model** section of STAR Methods),

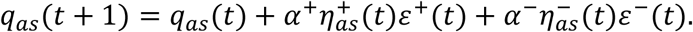

where *ε*^+^(*t*) ≡ max(0, *ε*(*t*)) and *ε*^−^(*t*) ≡ min(0, *ε*(*t*)). In the further downstream, the exploration and exploitation parameter *β* exists in the action choice probability,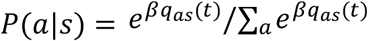. However, the separation for the sign of prediction error is already merged into the common policy parameter *q*_*as*_(*t*). In order to assume the imbalance of *β*, the policy parameter should be separated into 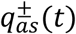, but it will cause divergence of the policy parameters, 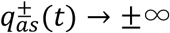. Thus, we consider only the imbalances of the learning rate and the trace factor for positive and negative prediction errors.

We compare the effects of imbalance in the learning rates and eligibility traces in the steady state of learning,

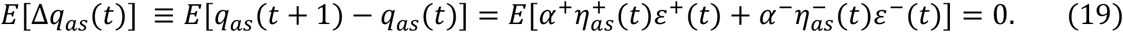

The eligibility traces 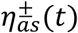 consist of the current and the past components,

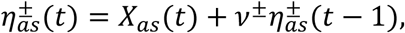

and Eq. (19) can be divided into the effects of immediate and delayed contingency of actions and outcomes,

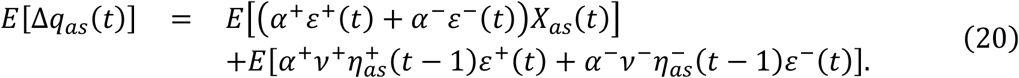

The imbalance of the learning rates *α*^+^≠ *α*^−^ disorders the contingency of immediate outcome, while the imbalance of the trace factors *ν*^+^≠ *ν*^−^ gives no effect in immediate contingency. In contrast, the imbalance of the trace factors *ν*^+^≠ *ν*^−^ causes temporally biased trace patterns 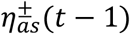 for past actions. Hence, the imbalance disorders the delayed contingency depending on the temporal correlation between the trace patterns and the outcome prediction error. Next, we discuss these differences in concrete situations.

#### 8. Imbalance effects in independent-trial condition

Previous works (Lefebvre *et al*., 2017) assuming the imbalance of learning rates have been discussed in tasks where the trials are temporally independent. In such cases, the difference between the imbalances of *α*^+^≠ *α*^−^ and *ν*^+^≠ *ν*^−^ is clear. Eq. (20) is calculated as

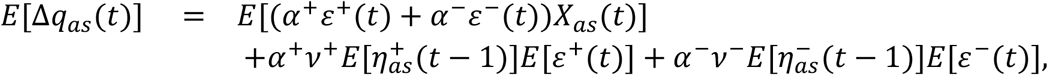

because the past eligibility traces 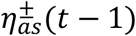 are independent of the current prediction errors *ε*^±^(*t*). The eligibility traces are described as 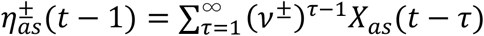 (see the **Separate eligibility trace model** section of STAR Methods), and the expectation of *X*_*a*_ does not depend on time in the steady state, *E*[*X*_*a*_(*t* − *τ*)] = *E*[*X*_*a*_]. Therefore, the above form is calculated as

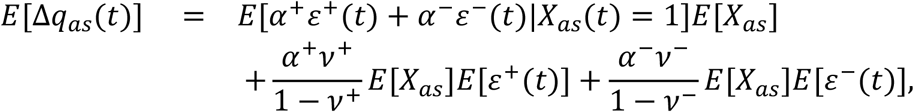

and the equilibrium equation *E*[Δ*q*_*as*_(*t*)] = 0 is obtained as

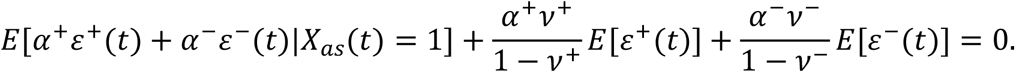

The past (last two) terms do not depend on state *s* and action *a*. Therefore, the past terms work as a uniform reinforcer *u*(*α*^±^, *ν*^±^),

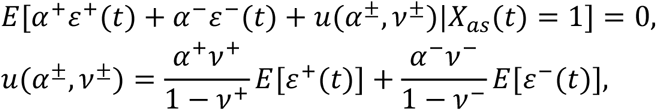

which never changes the value orders of policy parameters, whereas it can shift the values. The uniform reinforcer *u* is the function of the learning parameters, *α*^±^, *ν*^±^, and the averages of positive and negative prediction errors *E*[*ε*^±^(*t*)]. If the prediction error fluctuates, the positive and negative averages have non-zero values and can be distorted by the imbalance of the learning parameters.

The trace-factor imbalance affects only the uniform reinforcer *u*. Hence, the trace-factor imbalance can never change the choice preference in the steady state of learning.

In contrast, the learning-rate imbalance affects the update with immediate outcome contingency. This immediate effect is related to risk preference and aversion, which is included in the prospect theory (Kahneman and Tversky, 1979). The formulation in the reinforcement learning has been related to human behavior (Lefebvre *et al*., 2017).

Here we discussed risk preference and aversion, including trace-factor imbalance. Suppose safe and risky options. The safe option brings certain outcome. The risky option brings large or small outcomes probabilistically, and the average is equal to the outcome of the safe option. In this case, balanced reinforcement learning never causes bias in the choice preference, because the outcome expectations of safe and risky options are equal.

In the case of the learning-rate imbalance, the learning gain differs between positive and negative prediction errors (Supplementary Fig. 2a), in which the border of error zero corresponds to the “reference point” in prospect theory. The imbalance *α*^−^ < *α*^+^ leads to risk preference, and the imbalance *α*^−^ > *α*^+^ leads to risk aversion. Effect of the past component through the eligibility trace only results in upper (*α*^−^ < *α*^+^) or lower (*α*^−^ > *α*^+^) shift.

In contrast, the trace-factor imbalance causes only upper (*ν*^−^ < *ν*^+^) or lower (*ν*^−^ > *ν*^+^) shift whereas the learning gain is constant (Supplementary Fig. 2b).

## Supplemental items

**Supplementary Figure 1.**
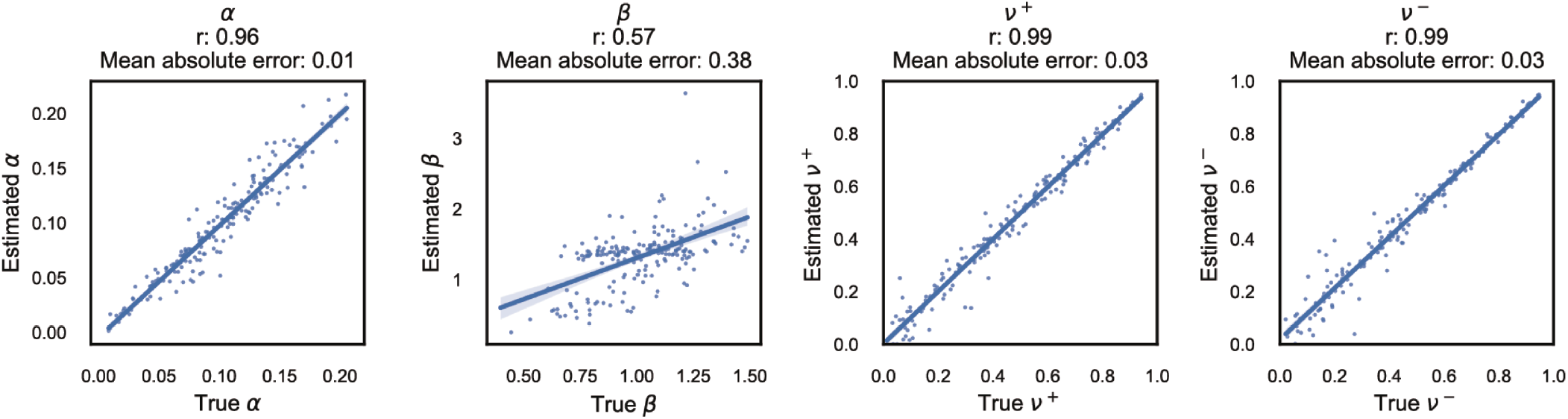
Parameter estimation, Related to STAR Methods. The horizontal and vertical axes represent the true and estimated parameters, respectively. The line and colored areas are the regression line and the 95% bootstrapped confidence interval, respectively.

**Supplementary Figure 2.**
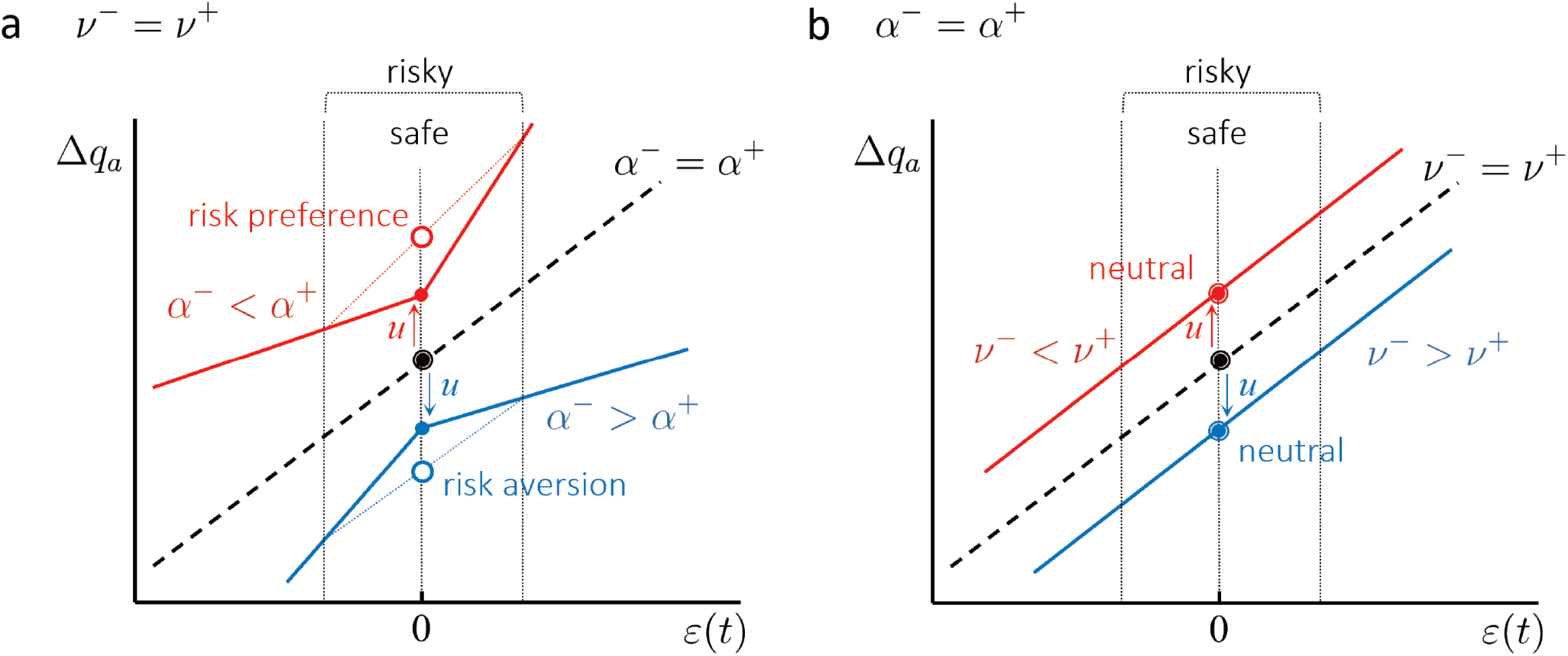
The difference between learning-rate and trace-factor imbalance in independent trial condition, Related to Figure 1. Updates of policy parameters depending on outcome prediction errors are plotted for the prediction error. (a) Learning updates under the learning-rate imbalance. The risk preference occurs because of the positive curvature (red lines; *α*^+^ > *α*^−^) around the expectation of mean outcomes that corresponds to the reference point in the prospect theory. In contrast, the negative curvature (blue lines; *α*^+^ < *α*^−^) causes risk aversion. The effect of eligibility trace causes a shift (*u*) in the update of policy parameters (see the theoretical derivation section of STAR Methods). The behavior depends only on the difference between policy parameters. Therefore, uniform shift does not cause any behavioral changes. (b) Learning updates under the trace-factor imbalance. The trace-factor imbalance cannot change the curvature. The behavior is invariant even if a shift (*u*) exists.

**Supplementary Table 1.**
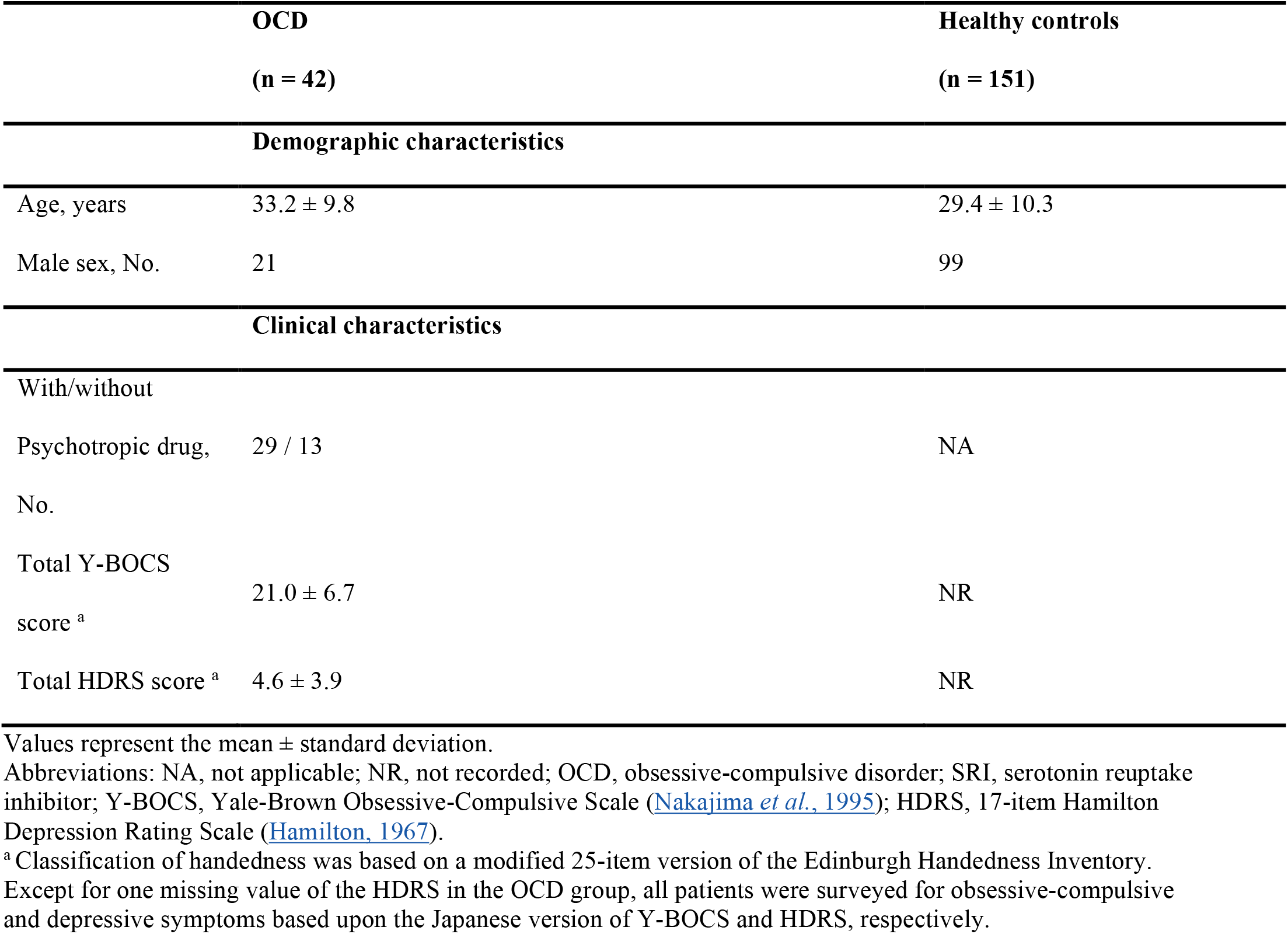
Participant demographic information, Related to STAR Methods. All demographic distributions are matched among groups (p > 0.05).

**Supplementary Table 2.** SRI doses and imipramine equivalent doses of each patient (n = 29), Related to STAR Methods.

| **SRI dose** | **Imipramine equivalent dose** |
| --- | --- |
| Paroxetine 5 mg | 18.75 mg |
| Paroxetine 15 mg | 56.25 mg |
| Paroxetine 20 mg | 75 mg |
| Paroxetine 25 mg | 93.75 mg |
| Paroxetine 30 mg | 112.5 mg |
| Paroxetine 30 mg | 112.5 mg |
| Escitalopram 10 mg | 75 mg |
| Escitalopram 10 mg | 75 mg |
| Escitalopram 10 mg | 75 mg |
| Clomipramine 50 mg | 62.5 mg |
| Paroxetine 40 mg | 150 mg |
| Paroxetine 40 mg | 150 mg |
| Paroxetine 50 mg | 187.5 mg |
| Paroxetine 50 mg | 187.5 mg |
| Paroxetine 50 mg | 187.5 mg |
| Fluvoxamine 150 mg | 150 mg |
| Fluvoxamine 150 mg | 150 mg |
| Fluvoxamine 150 mg | 150 mg |
| Sertraline 75 mg + Escitalopram 10 mg | 187.5 mg |
| Clomipramine 150 mg | 187.5 mg |
| Clomipramine 75 mg | 93.75 mg |
| Escitalopram 10 mg | 75 mg |
| Fluvoxamine 100 mg | 100 mg |
| Paroxetine 30 mg | 112.5 mg |
| Paroxetine 50 mg | 187.5 mg |
| Escitalopram 20 mg | 150 mg |
| Paroxetine 50 mg | 187.5 mg |
| Paroxetine 50 mg | 187.5 mg |
| Paroxetine 30 mg | 112.5 mg |

## References

Abe, Y., Sakai, Y., Nishida, S., Nakamae, T., Yamada, K., Fukui, K., and Narumoto, J. (2015). Hyper-influence of the orbitofrontal cortex over the ventral striatum in obsessive-compulsive disorder. Eur Neuropsychopharmacol 25, 1898–1905. 10.1016/j.euroneuro.2015.08.017.

Anderson, M.J., Walsh, D.C.I., Robert Clarke, K., Gorley, R.N., and Guerra-Castro, E. (2017). Some solutions to the multivariate Behrens-Fisher problem for dissimilarity-based analyses. Australian & New Zealand Journal of Statistics 59, 57–79. 10.1111/anzs.12176.

Banca, P., Vestergaard, M.D., Rankov, V., Baek, K., Mitchell, S., Lapa, T., Castelo-Branco, M., and Voon, V. (2014). Evidence Accumulation in Obsessive-Compulsive Disorder: the Role of Uncertainty and Monetary Reward on Perceptual Decision-Making Thresholds. Neuropsychopharmacology. 10.1038/npp.2014.303.

Bergstra, J.S., Bardenet, R., Bengio, Y., and Kégl, B. (2011). Algorithms for hyper-parameter optimization. pp. 2546–2554.

Brzosko, Z., Mierau, S.B., and Paulsen, O. (2019). Neuromodulation of Spike-Timing-Dependent Plasticity: Past, Present, and Future. Neuron 103, 563–581. 10.1016/j.neuron.2019.05.041.

Brzosko, Z., Schultz, W., and Paulsen, O. (2015). Retroactive modulation of spike timing-dependent plasticity by dopamine. eLife 4, e09685. 10.7554/eLife.09685.

Brzosko, Z., Zannone, S., Schultz, W., Clopath, C., and Paulsen, O. (2017). Sequential neuromodulation of Hebbian plasticity offers mechanism for effective reward-based navigation. eLife 6, e27756. 10.7554/eLife.27756.

Campello, R.J., Moulavi, D., Zimek, A., and Sander, J. (2015). Hierarchical density estimates for data clustering, visualization, and outlier detection. ACM Transactions on Knowledge Discovery from Data (TKDD) 10, 5.

Cavaccini, A., Gritti, M., Giorgi, A., Locarno, A., Heck, N., Migliarini, S., Bertero, A., Mereu, M., Margiani, G., Trusel, M., et al. (2018). Serotonergic Signaling Controls Input-Specific Synaptic Plasticity at Striatal Circuits. Neuron 98, 801–816 e807. 10.1016/j.neuron.2018.04.008.

Cavedini, P., Gorini, A., and Bellodi, L. (2006). Understanding obsessive-compulsive disorder: focus on decision making. Neuropsychol Rev 16, 3–15. 10.1007/s1065-006-9001-y.

Chamberlain, S.R., and Menzies, L. (2009). Endophenotypes of obsessive-compulsive disorder: rationale, evidence and future potential. Expert Rev Neurother 9, 1133–1146. 10.1586/ern.09.36.

Cleeremans, A. (2009). Implicit Learning and Implicit Memory. In Encyclopedia of Consciousness, W.P. Banks, ed. (Academic Press), pp. 369–381. https://doi.org/10.1016/B978-012373873-8.00047-5.

Daw, N.D. (2011). Trial-by-trial data analysis using computational models. Decision making, affect, and learning: Attention and performance XXIII 23, 3–38.

First, M.B., Spizer, R.L., Gibbon, M., and Williams, J.B.W. (1994). Structures Clinical Interview for Axis I DSM-IV Disorders-Patient Edition (CID-I/P). New York: Biometrics Research Department, New York State Psychiatric Institute.

Gerstner, W., Lehmann, M., Liakoni, V., Corneil, D., and Brea, J. (2018). Eligibility Traces and Plasticity on Behavioral Time Scales: Experimental Support of NeoHebbian Three-Factor Learning Rules. Frontiers in neural circuits 12, 53. 10.3389/fncir.2018.00053.

Gillan, C.M., Kalanthroff, E., Evans, M., Weingarden, H.M., Jacoby, R.J., Gershkovich, M., Snorrason, I., Campeas, R., Cervoni, C., and Crimarco, N.C. (2019). Comparison of the Association Between Goal-Directed Planning and Self-reported Compulsivity vs Obsessive-Compulsive Disorder Diagnosis. JAMA psychiatry, 1–10.

Gillan, C.M., and Robbins, T.W. (2014). Goal-directed learning and obsessive-compulsive disorder. Philos Trans R Soc Lond B Biol Sci 369. 10.1098/rstb.2013.0475.

Gillan, C.M., and Sahakian, B.J. (2015). Which is the driver, the obsessions or the compulsions, in OCD? Neuropsychopharmacology 40, 247–248. 10.1038/npp.2014.201.

Graybiel, A.M. (2000). The basal ganglia. Curr Biol 10, R509–511. 10.1016/s0960-9822(00)00593-5.

Hamilton, M. (1967). Development of a rating scale for primary depressive illness. The British journal of social and clinical psychology 6, 278–296. 10.1111/j.2044-8260.1967.tb00530.x.

Hauser, T.U., Moutoussis, M., Iannaccone, R., Brem, S., Walitza, S., Drechsler, R., Dayan, P., and Dolan, R.J. (2017). Increased decision thresholds enhance information gathering performance in juvenile Obsessive-Compulsive Disorder (OCD). PLoS Comput Biol 13, e1005440. 10.1371/journal.pcbi.1005440.

He, K., Huertas, M., Hong, S.Z., Tie, X., Hell, J.W., Shouval, H., and Kirkwood, A. (2015). Distinct Eligibility Traces for LTP and LTD in Cortical Synapses. Neuron 88, 528–538. 10.1016/j.neuron.2015.09.037.

Inada, T., and Inagaki, A. (2015). Psychotropic dose equivalence in Japan. Psychiatry Clin Neurosci 69, 440–447. 10.1111/pcn.12275.

Izhikevich, E.M. (2007). Solving the distal reward problem through linkage of STDP and dopamine signaling. Cereb Cortex 17, 2443–2452. 10.1093/cercor/bhl152.

Kahneman, D., and Tversky, A. (1979). Prospect Theory: An Analysis of Decision under Risk. Econometrica 47, 263–291. 10.2307/1914185.

Koran, L.M., Hanna, G.L., Hollander, E., Nestadt, G., Simpson, H.B., and American Psychiatric, A. (2007). Practice guideline for the treatment of patients with obsessive-compulsive disorder. Am J Psychiatry 164, 5–53.

Kravitz, A.V., Tye, L.D., and Kreitzer, A.C. (2012). Distinct roles for direct and indirect pathway striatal neurons in reinforcement. Nat Neurosci 15, 816–818. 10.1038/nn.310.

Lefebvre, G., Lebreton, M., Meyniel, F., Bourgeois-Gironde, S., and Palminteri, S. (2017). Behavioural and neural characterization of optimistic reinforcement learning. Nature Human Behaviour 1, 067.

Nakajima, T., Nakamura, M., Taga, C., Yamagami, S., Kiriike, N., Nagata, T., Saitoh, M., Kinoshita, T., Okajima, Y., Hanada, M., and et al. (1995). Reliability and validity of the Japanese version of the Yale-Brown Obsessive-Compulsive Scale. Psychiatry Clin Neurosci 49, 121–126. 10.1111/j.1440-1819.1995.tb01875.x.

Noguchi, K., Gel, Y.R., Brunner, E., and Konietschke, F. (2012). nparLD: an R software package for the nonparametric analysis of longitudinal data in factorial experiments. Journal of Statistical software 50.

Nonomura, S., Nishizawa, K., Sakai, Y., Kawaguchi, Y., Kato, S., Uchigashima, M., Watanabe, M., Yamanaka, K., Enomoto, K., and Chiken, S. (2018). Monitoring and updating of action selection for goal-directed behavior through the striatal direct and indirect pathways. Neuron 99, 1302-1314.e1305.

Palminteri, S., and Lebreton, M. (2022). The computational roots of positivity and confirmation biases in reinforcement learning. Trends in Cognitive Sciences 26, 607–621. https://doi.org/10.1016/j.tics.2022.04.05.

Pauls, D.L., Abramovitch, A., Rauch, S.L., and Geller, D.A. (2014). Obsessive-compulsive disorder: an integrative genetic and neurobiological perspective. Nat Rev Neurosci 15, 410–424. 10.1038/nrn3746.

Sakai, Y., Narumoto, J., Nishida, S., Nakamae, T., Yamada, K., Nishimura, T., and Fukui, K. (2011). Corticostriatal functional connectivity in non-medicated patients with obsessive-compulsive disorder. Eur Psychiatry 26, 463–469. 10.1016/j.eurpsy.2010.09.005.

Salkovskis, P.M. (1999). Understanding and treating obsessive-compulsive disorder. Behav Res Ther 37 Suppl 1, S29–52.

Schultz, W., Dayan, P., and Montague, P.R. (1997). A neural substrate of prediction and reward. Science 275, 1593–1599. 10.126/science.275.5306.1593.

Shen, W., Flajolet, M., Greengard, P., and Surmeier, D.J. (2008). Dichotomous dopaminergic control of striatal synaptic plasticity. Science 321, 848–851. 10.1126/science.1160575.

Sugiura, Y., and Tanno, Y. (2000). Self-report inventory of obsessive-compulsive symptoms: Reliability and validity of the Japanese version of the Padua Inventory.

Sutton, R.S., and Barto, A.G. (1998). Reinforcement Learning: An Introduction. MIT Press, Cambridge, MA.

Tanaka, S.C., Shishida, K., Schweighofer, N., Okamoto, Y., Yamawaki, S., and Doya, K. (2009). Serotonin affects association of aversive outcomes to past actions. J Neurosci 29, 15669–15674. 10.1523/JNEUROSCI.2799-09.209.

Vaghi, M.M., Luyckx, F., Sule, A., Fineberg, N.A., Robbins, T.W., and De Martino, B. (2017). Compulsivity Reveals a Novel Dissociation between Action and Confidence. Neuron.

Wakabayashi, A., Tojo, Y., Baron-Cohen, S., and Wheelwright, S. (2004). [The Autism-Spectrum Quotient (AQ) Japanese version: evidence from high-functioning clinical group and normal adults]. Shinrigaku Kenkyu 75, 78–84. 10.4992/jjpsy.75.78.

Wilson, R.C., and Collins, A.G.E. (2019). Ten simple rules for the computational modeling of behavioral data. eLife 8, e49547. 10.7554/eLife.49547.

Yagishita, S., Hayashi-Takagi, A., Ellis-Davies, G.C., Urakubo, H., Ishii, S., and Kasai, H. (2014). A critical time window for dopamine actions on the structural plasticity of dendritic spines. Science 345, 1616–1620. 10.1126/science.1255514.

Yazdani, S., Vahabie, A.-H., Araabi, B.N., and Ahmadabadi, M.N. (2018). Better than maximum likelihood estimation of model-based and model-free learning style. bioRxiv, 296335.

